# Unraveling the Complexity of Amyloid Polymorphism Using Gold Nanoparticles and Cryo-EM

**DOI:** 10.1101/754846

**Authors:** Urszula Cendrowska, Paulo Jacob Silva, Nadine Ait-Bouziad, Marie Müller, Zekiye Pelin Guven, Sophie Vieweg, Anass Chiki, Lynn Radamaker, Senthil Thangaraj, Marcus Fändrich, Francesco Tavanti, Maria Cristina Menziani, Alfredo Alexander-Katz, Francesco Stellacci, Hilal A. Lashuel

**Author notes:** These authors contributed equally (Urszula Cendrowska, Paulo Jacob Silva). Corresponding Authors Hilal A. Lashuel, Francesco Stellacci. Significance Statement The ability of proteins to self-assemble into different types of fibrils with distinct morphologies has been linked to the pathological and clinical heterogeneity of amyloid diseases such as Alzheimer’s and Parkinson’s disease. Herein, we describe novel nanoparticles (NPs) that efficiently label amyloid fibrils produced in vitro or isolated from postmortem tissues, under hydrating conditions and in such a way to unmask their polymorphism and morphological features. Using these NPs, we show that pathological aggregates exhibit exceptional morphological homogeneity compared to amyloid fibrils produced in vitro, consistent with the emerging view that the physiologic milieu is a key determinant of amyloid fibril strains. These advances pave the way for elucidating the structural basis of amyloid strains and toxicity.

## Abstract

The misfolding and self-assembly of proteins into β-sheet-rich amyloid fibrils of various structures and morphologies is a hallmark of several neurodegenerative and systemic diseases. Increasing evidence suggests that amyloid polymorphism gives rise to different strains of amyloids with distinct toxicity and pathology-spreading properties. Validating this hypothesis is challenging due to a lack of tools and methods that allow for the direct characterization of amyloid polymorphism in hydrated and complex biological samples. Here, we report on the use of 11-mercapto-1-undecanesulfonate-coated gold nanoparticles (NPs) to label the edges of synthetic, recombinant and native amyloid fibrils to assess amyloid morphological polymorphism using cryogenic transmission electron microscopy (cryo-EM). The fibrils studied were derived from amyloid proteins involved in disorders of the central nervous system (amyloid-β, tau, α-synuclein) and in systemic amyloidosis (a fragment of an immunoglobulin λ light chain). The labeling efficiency enabled imaging and characterization of amyloid fibrils of different morphologies under hydrated conditions using cryo-EM. These NPs allowed for the visualization of morphological features that are not directly observed using standard imaging techniques, including TEM with use of the negative stain or cryo-EM imaging. We also demonstrate the use of these NPs to label native paired helical filaments (PHFs) from the postmortem brain of an Alzheimer’s disease patient, as well as amyloid fibrils extracted from the heart tissue of a patient suffering from systemic amyloid light-chain (AL) amyloidosis. Analysis of the cryo-EM images of amyloids decorated with NPs shows exceptional homogeneity across the fibrils derived from human tissue in comparison to fibrils aggregated in vitro. The use of these NPs enabled us to gain novel insight into the structural features that distinguish amyloid fibrils formed in vivo from those formed in cell-free in vitro systems. Our findings demonstrate that these NPs represent a powerful tool for rapid imaging and profiling of amyloid morphological polymorphism in different types of samples, including those derived from complex biological aggregates found in human tissue and animal models of amyloid diseases. These advances should not only facilitate the profiling and characterization of amyloids for structural studies by cryo-EM but also pave the way to elucidate the structural basis of amyloid strains and toxicity and possibly the correlation between the pathological and clinical heterogeneity of amyloid diseases.

---

Amyloids are insoluble β-sheet-rich fibrillar protein aggregates found in pathological deposits and inclusions that characterize several neurodegenerative and systemic diseases (1). These diseases span from degenerative diseases of the central nervous system, such as Alzheimer’s disease (AD) and Parkinson’s disease (PD), to systemic disorders, such as type II diabetes and systemic amyloidosis (1–5). Although the mechanisms of amyloid formation in vivo and the nature of the toxic species remain subjects of intense investigation and debate, there is converging evidence that the fibrillization process plays a central role in neurodegeneration and pathology spreading in neurodegenerative diseases. Therefore, the prevention or inhibition of protein misfolding, aggregation and pathology spreading remains the most actively pursued goal to develop therapeutic strategies against amyloid diseases.

Increasing evidence from in vitro and animal models as well as structural studies on both ex vivo and synthetic aggregates have shown that amyloid-forming proteins can self-assemble into fibrils of distinct morphologies and toxic properties. Several studies have shown that amyloidogenic proteins such as amyloid-β (Aβ), α-synuclein and tau protein self-assemble in vitro into fibrils of different morphologies, e.g., ribbon, helical and straight fibrils, depending on their growth conditions (6–10). Even in the same sample, different types of fibril morphologies have been shown to form and coexist, suggesting that they are either in an equilibrium or are derived from morphologically distinct intermediates on the pathway to aggregation (11, 12). The polymorphism of cross-β fibrils has also been observed in ex vivo fibrils, for instance isolated tau fibrils from the brain of patients with AD (13) or amyloid fibrils from patients and animals with systemic amyloidosis of amyloid A (AA), transthyretin (ATTR) and light chain (AL) (14).

Recent studies have demonstrated that introducing recombinant preformed fibrils or amyloid aggregates derived from diseased brains into neurons induces amyloid aggregation. The same process was observed when these fibrils were introduced directly to a specific region of the animal model brain. In these seeding-based models, distinct fibrillar polymorphs lead to different patterns of pathology spreading and/or toxicity (15–18). Tycko and coworkers have developed a method that enables the amplification of native amyloids by seeding monomeric Aβ with different fibril types isolated from different AD patients or brain regions (19, 20). Their findings suggest a correlation between the amyloid fibril molecular structure, amyloid toxicity and symptomatology. Peng and colleagues (16) demonstrated that isolated cytoplasmic brain aggregates from Parkinson’s disease (Lewy bodies, LBs) and multiple system atrophy (MSA; glial inclusions, GCIs) exhibit distinct conformations and drastically different seeding capacities of α-synuclein. GCI α-synuclein forms structures that are more compact and approximately a thousand-fold more potent than those of LB α-synuclein in terms of seeding capacity. Similar observations were made with tau (21), where tau isolated from five different tauopathies was shown to indefinitely propagate distinct amyloid conformations in a clonal fashion when inoculated into a cellular model. These findings and the availability of cellular and animal models of amyloid propagation provide a unique opportunity to investigate the relationship between amyloid molecular and morphological polymorphism, neurodegeneration and pathology spreading in neurodegenerative diseases and are important for developing effective therapeutics. Nonetheless, it remains challenging to elucidate the structural basis and role of fibril polymorphism in human-derived material due to the minute amount of these aggregates and the complexity of the samples.

Amyloid polymorphism can occur at different levels. Tycko and coworkers have established that the same protein can have different molecular arrangements when forming an amyloid fibril, and in this paper, we refer to this as molecular polymorphism (22). Mezzenga and coworkers have shown that the shape, e.g., the symmetry and thickness of a fibril, varies depending on the number of protofilaments that compose the fibril (23). We refer to this as morphological polymorphism. These two types of polymorphism are not mutually exclusive and could coexist.

Molecular polymorphism has been investigated using solid-state NMR (ssNMR) spectroscopy, microcrystallography and X-ray diffraction, while transmission electron microscopy (TEM), scanning probe microscopy and atomic force microscopy (AFM) have been used to investigate morphological polymorphism (24, 25). These techniques have similar limitations: 1) they are time consuming; 2) they require large amounts of protein (e.g., ssNMR); 3) they can be applied only to small amyloid-forming peptides (e.g., microcrystallography); 4) they do not allow imaging of amyloids in complex systems under native and hydrated conditions (e.g., TEM); or 5) they require the presence of a substrate that may strongly influence the aggregation pathway of the growing fibrils (e.g., AFM) (26). For example, direct investigation of the molecular basis of amyloid polymorphism of Aβ in AD brain tissue using ssNMR spectroscopy was only possible by seeding brain-derived aggregates (seeds) to an excess of isotopically labeled monomeric proteins, yielding hundreds of milligrams of fibrils (27–29). In TEM, samples are dried on the grid and stained with an electron-dense staining agent before imaging. Although this technique allows for rapid characterization of the structural morphology of the fibrils (30), it may introduce several artifacts, such as flattening of the structure or incomplete stain embedding (31). Cryo-EM can address these shortcomings because it does not require drying, staining or surface deposition. With this technique, the sample is frozen in a thin layer of vitreous ice, which allows fibrils to be imaged in their hydrated state. This method is increasingly used to investigate the structure of amyloid fibrils prepared in vitro or derived from different tissues (11, 13, 32). However, the measurements are based on the weak contrast between fibrils and glassy ice and require a large number of images and advanced data analysis techniques to extract morphological information.

Herein, we describe the synthesis and application of gold nanoparticles (NPs) that efficiently label amyloid fibrils in a specific manner and in such a way so to unmask their polymorphism and morphological features. Our NPs combined with cryo-EM enable rapid screening and detailed characterization of amyloid morphological polymorphism under hydrated conditions Importantly, the resulting information cannot be directly observed using standard imaging techniques. One of the unique features of these NPs is that they efficiently label amyloid fibrils produced from a wide range of disease-associated recombinant proteins or synthetic peptides in vitro ((e.g. amyloid beta peptides, alpha-synuclein, TDP-43, Tau, immunoglobulin λ light chain and exon1 of the Huntingtin protein), despite the diversity of their amino acid sequence, secondary structure contents, and fibrillar morphology. Furthermore, our NPs efficiently label ex vivo disease-associated fibrils isolated from pathological aggregates derived from the brain and heart of patients that died from Alzheimer’s disease and AL amyloidosis, respectively. Our results also show that brain and heart-derived amyloid fibrils isolated from pathological aggregates exhibit exceptional morphological homogeneity, in terms of their helical periodicity, in comparison to amyloid fibrils produced from recombinant proteins or peptide in vitro. These findings are consistent with and strongly support the emerging view that the physiologic milieu is a key determinant of amyloid fibril strains.

## Results

### MUS:OT NPs successfully label different polymorphs of amyloid fibrils

Noble metal NPs have been previously used to study amyloid fibrils. Since 1971, gold NPs conjugated with a specific antibody have been successfully used to label amyloids (33). This technique, called immunogold labeling, allows for the localization of amyloids in tissue sections and characterization of the sequence and structural properties of fibrils in solution (34, 35). However, with this technique, NPs do not anchor directly on the fibrils but are located as far as 30 nm away due to the presence of the bulky antibody and linker. Therefore, the NPs used under these conditions cannot provide information about the structural or morphological features of amyloid fibrils, which renders this technique unsuitable for the characterization of fibril polymorphism (36). There are also examples of amyloids being labeled using a nanomaterial without spacers, such as maghemite NPs (37, 38), gold nanorods (39) or gold NPs (40, 41). Due to their bulky size and the nonspecific nature of their interactions with amyloid fibrils, these nanomaterials are also unsuitable for unravelling morphological features and heterogeneity of fibrils.

To address these limitations, we sought to develop new tools that leverage the advantages offered by NPs, e.g., small size with significant electron density, to enable determining amyloid morphological polymorphism under hydrated conditions using cryo-EM. Towards this goal, we first synthesized NPs with various charges and ligands: positively charged (*N,N,N-trimethyl(11-mercaptoundecyl)ammonium chloride,* TMA), negatively charged (*11-mercapto-1-undecanesulfonate,* MUS, *11-mercaptoundecylphosphoric acid,* MUP), mixed ligand negatively charged (mixtures of MUS and 1-octanethiol referred to as MUS:OT A and MUS:OT B) and zwitterionic NPs (*3-[(11-mercapto-undecyl)-N,N-dimethylamino]propane-1-sulfonate,* ZW). Positively charged and zwitterionic NPs were synthesized via modification of the method developed by Zheng et al. (42), while anionic NPs were synthesized via a one-phase method, as described previously (42) and in the Supporting Information. The ligand shell composition was determined by ^1^H-NMR spectroscopy after etching the gold core with iodine (43), while the size distribution of the Au core was determined using TEM (Fig. 1A – B, Supporting Information, and Fig. S1).

**Fig. 1.**
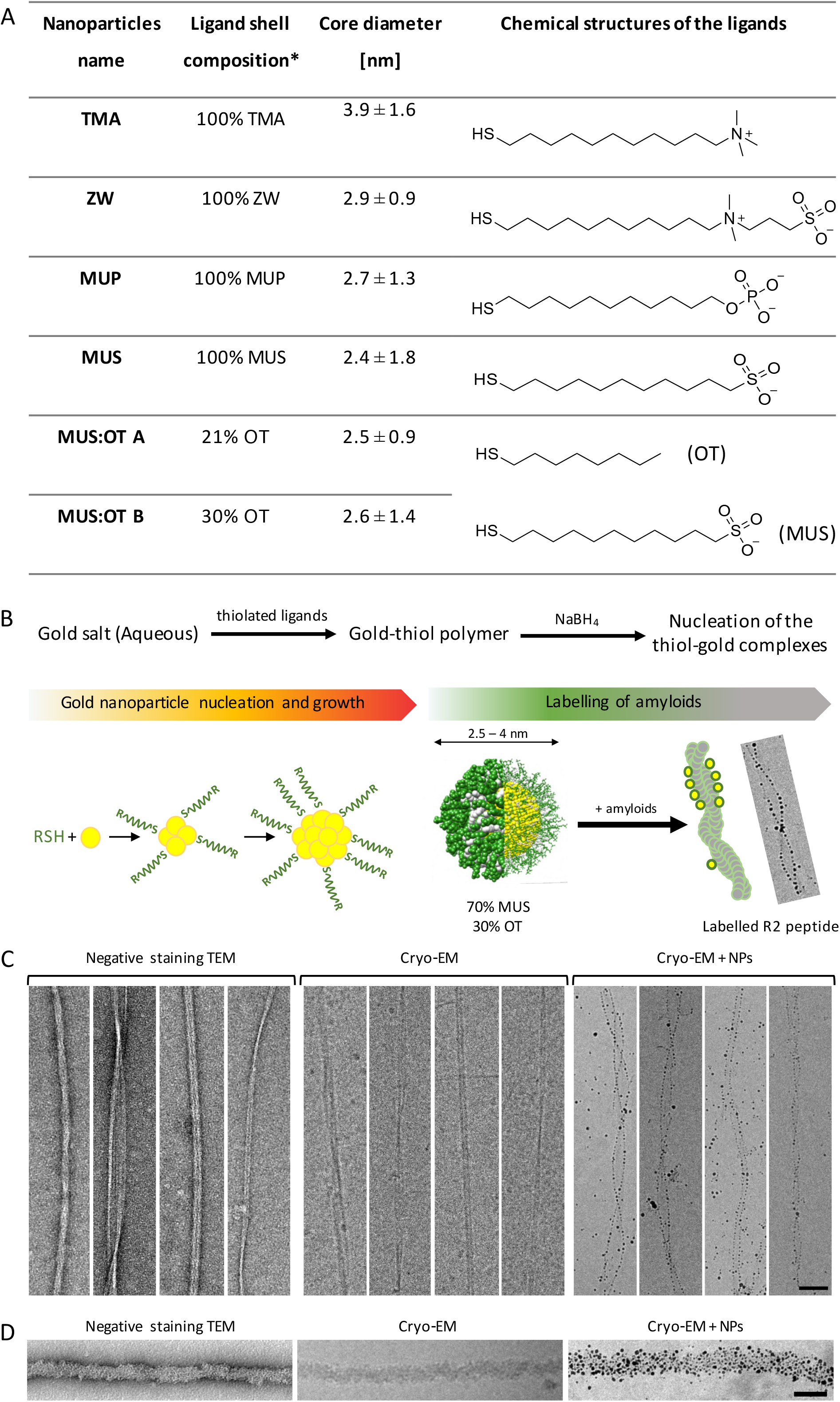
A) Table presenting the NPs and their characteristics – ligand shell composition, core size and chemical structure of the protective ligands. B) Simplified scheme of NP synthesis with the cartoon depicting the nucleation and the amyloid-labeling process. NPs are synthesized from gold salt upon addition of the chosen ligand(s), which triggers the creation of a gold-thiolate complex. A reducing agent is added dropwise to the gold-thiol mixture at the end of the process and causes the nucleation of the NPs, which subsequently grow to their final size that spans from 1 to 5 nm (mathematical simulation of MUS:OT B NPs, where the gold core is depicted in yellow, the MUS ligand in green and the OT ligand in white). Amyloid fibrils become decorated through the adsorption of NPs on their surface. C) Comparison of R2 fibrils imaged by negative staining, cryo-EM and cryo-EM images of the fibrils decorated with MUS:OT A NPs. The different morphologies of the fibrils, including straight and twisted fibrils, are clearly highlighted by the NPs. D) Comparison of TDP-43 fibrils imaged by negative staining, cryo-EM and cryo-EM images of the fibrils decorated with MUS:OT A NPs. Unlike other amyloids, TDP-43 fibrils do not reveal helical morphology yet still become evenly decorated. Scale bars are 50 nm. *Ligand ratio calculated from ^1^H NMR analysis after decomposition of the core.

To achieve efficient labeling and visualization, we used gold NPs with a core diameter of ∼3 nm, as smaller particles are difficult to visualize in conventional cryo-EM and larger particles may result in a lower morphological accuracy and spatial localization of the fibril. To determine ligands with the best labelling efficiency, we first examined the interaction between Aβ_40_ fibrils and the different types of gold NPs by coincubating fibrils with different NPs and then imaging them using cryo-EM. The Aβ_40_ peptide is one of the constituents of amyloid plaques and one of the key defining pathological hallmarks of AD, together with the protein tau. Previous studies have shown that Aβ_40_ self-assembles and forms fibrils of different morphologies under different conditions (7, 11), which makes it a good model system to study how our NPs can be used to characterize fibril polymorphs. Among all of the synthesized NPs, the mixture of OT and MUS ligands on the gold surface was the most efficient in labeling Aβ_40_ fibrils, as the other NPs tested either did not attach efficiently to the fibril surface or formed large fibril-NP aggregates (Supporting Information, Fig. S2). We observed that both types of MUS:OT NPs showed similar amyloid labeling efficiency despite the small difference in the hydrophobic ligand ratio (Fig. 2). Higher ratios of OT in the ligand shell often resulted in reduced NP solubility under the buffer conditions used in this study. These observations combined with molecular dynamics simulations studies (Supporting information) lead us to hypothesize that labeling efficiency depends on the hydrophobic contacts between the NPs and the amyloid fibrils.

**Fig. 2.**
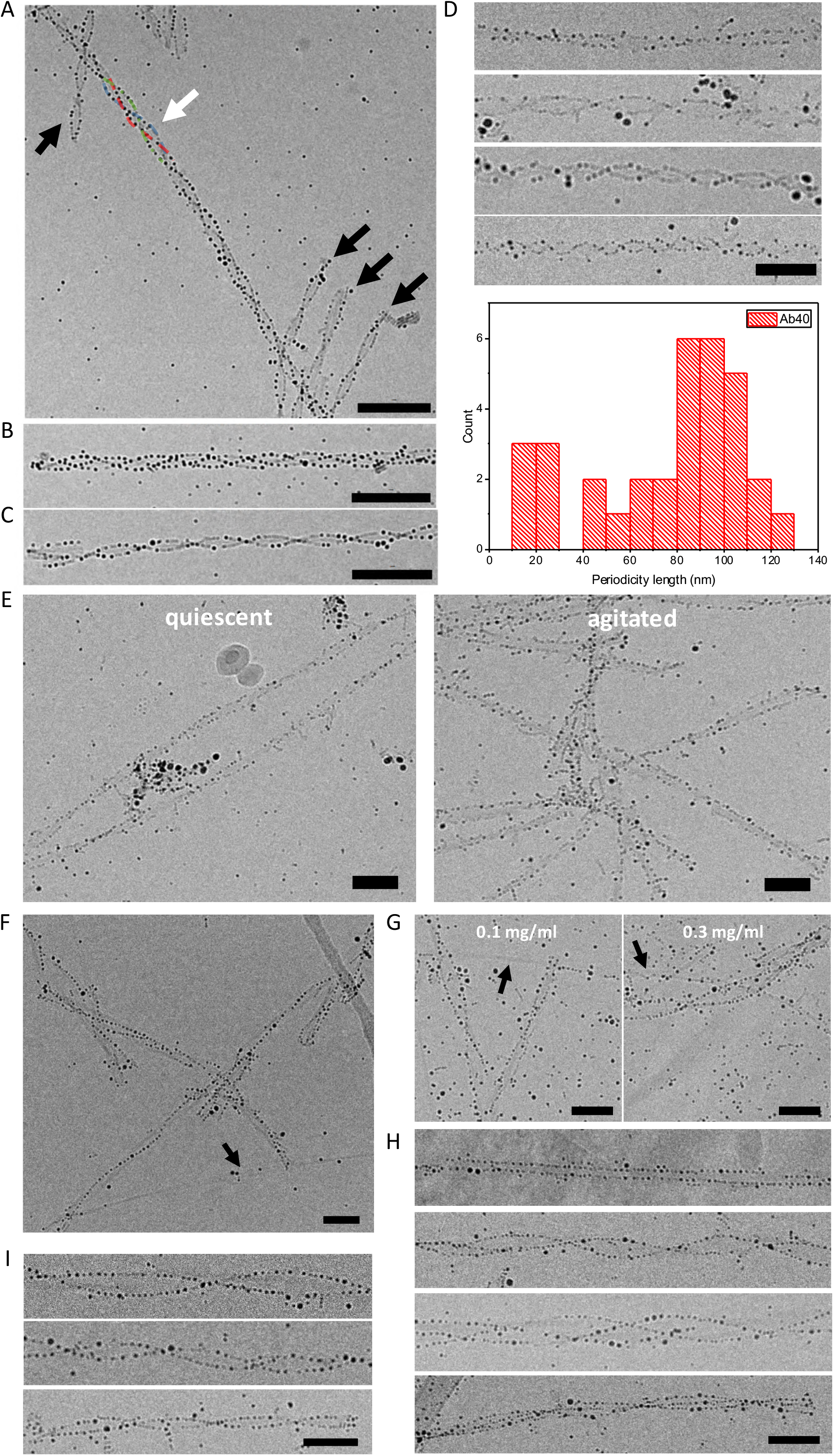
MUS:OT NPs decorating Aβ_40_ and R2 fibrils. A) Mixture of three-fold and two-fold Aβ_40_ fibrils within a sample detected with the use of MUS:OT NPs. Black arrows point to two-fold fibrils, and the white arrow points to a three-fold symmetric fibril. B) and C) show higher magnification images of three- and two-fold symmetric fibrils, respectively. D) Quantification of the two-fold fibril periodicity distribution within a sample. Upper panel: micrographs of Aβ_40_ amyloid fibrils with various periodicity. Lower panel: periodicity distribution of two-fold fibrils, highlighting the diversity of Aβ_40_ fibril morphological polymorphs in a single sample. E) Comparison between fibrils grown under quiescent conditions decorated by gold NPs unraveling their twist and short, ribbon-like shaped fibrils that were agitated during the aggregation process. F) Decorated twisted fibrils coexist with bare and thin R2 fibrils. The black arrow points at the bare fibril. G) Undecorated thin fibrils visible in the sample with increased concentration of NPs. Black arrows point to the undecorated amyloid fibrils, which show a specific morphology. H) Different morphologies of the R2 fibrils are easily discriminated owing to the labeling with NPs. I) Twisted R2 fibrils of different periodicities. Micrographs shown on the A), B) and C) panels contain amyloids decorated with the use of MUS:OT B NPs, while the rest of the panels show fibrils covered with MUS:OT A NPs. All scale bars are 50 nm.

We also examined the influence of our nanomaterial on the secondary structure and fibril morphology of Aβ_40_ by coincubating Aβ_40_ with NPs during the aggregation process and on the mature fibrils obtained at the end of the aggregation reaction. As shown in Fig. S3, the spectra of the fibrils incubated with MUS:OT A NPs (both during the aggregation process and of mature fibrils) are comparable to those of the control sample (i.e., CD spectra of fibrils incubated in the absence of NPs) and did not change significantly over time. This result is a strong indication that the presence of NPs does not modify the secondary structure of the amyloid fibrils and that the possible influence on the fibril structure upon decoration with NPs is minimal.

To illustrate the advantage of using our NPs, we imaged fibrils derived from the second repeat region of tau via negative staining TEM, cryo-EM and cryo-EM after coincubation with NPs and fibrils. Fig. 1C can be used as a comparison between imaging fibrils with the use of all three techniques. As shown in Fig 1C, NPs facilitate the visualization of the amyloid edges in cryo-EM while retaining the main advantage of cryo-EM relative to that of negative stain TEM, i.e., eliminating the need for drying.

To determine whether our NPs could decorate fibrils that do not present clear twisted morphologies, we incubated NPs with transactive response DNA-binding protein 43 (TDP-43) fibrils, which is implicated in several neurodegenerative conditions including amyotrophic lateral sclerosis (ALS), subsets of frontotemporal dementia (FTLD-TDP) and AD (44). We were able to obtain uniform decoration with MUS:OT A NPs across the entirety of the fibrils (Fig. 1D), which significantly facilitated the detection of the fibrils in the ice. These findings demonstrate that our NPs enable fast characterization of fibril morphological polymorphism in in vitro samples of different amyloids and reveal structural features that are not readily visible by standard cryo-EM in the absence of extensive image analysis.

### MUS:OT NPs enable rapid profiling and characterization of the polymorphism of the AD associated Aβ_40_ and tau-derived R2 peptides

MUS:OT NPs were used to perform an in-depth analysis of the polymorphism of Alzheimer’s associated Aβ_40_ and an amyloidogenic tau-derived peptide. Cryo-EM images of Aβ_40_ after 24 h of incubation with gold MUS:OT NPs under shaking conditions revealed homogenously decorated fibrils (Fig. 2A – E). The NPs are evenly distributed lining the edges of the fibrils, which increased their contrast under cryo-EM facilitating their detection on the grid. The fibril edges are easily discerned from the background, allowing for immediate determination of the type of morphological polymorph. We observe that Aβ_40_ fibrils exist predominantly as two classes of polymorphs: two- and three-folded helical symmetries around the longitudinal axis (Fig. 2A – C). The NP labeling facilitated the measurement of the crossover length distribution of the two-fold twisted fibril population (Fig. 2D), facilitating image analysis and quantification of the fibrils’ morphological parameters. Cryo-EM technique projects three-dimensional objects as a two-dimensional image. Two-fold symmetric images can be generated by the projection of twisted ribbons. More complex geometries may have the appearance of a three-fold symmetry that stems either from a twisted ribbon, a triangular cross-section, or helical ribbons (23). In this paper, we refer to the apparent image symmetry, not to the effective object symmetry (Fig. S4).

To investigate the effect of the aggregation conditions on the polymorphic population, we prepared Aβ_40_ fibrils using two different procedures: with and without agitation. The high affinity of the NPs to the fibril edges allows for rapid determination of the differences in the structural polymorph distribution between these two conditions. We observe that fibrils formed under quiescent conditions are longer and present a twisted helical symmetry, while those prepared under agitation are shorter and resemble straight ribbons devoid of any visible twist (Fig. 2E).

Tau is a natively unfolded protein, but under pathological conditions, it undergoes conformational changes that render the protein more prone to aggregation into paired helical filaments (PHFs). PHFs are the main component of the intracellular neurofibrillary tangles in the brains of AD patients (45). Tau’s ability to adopt β-sheet conformations, necessary for amyloid aggregation, relies mainly on small peptide motifs located in the second, third and fourth microtubule binding domains (MTBDs) of tau (46). To determine if the MUS:OT NPs could bind and label fibrils derived from other amyloid-forming peptides, we incubated them with fibril preparations of the MTBD-derived peptide (repeat unit R2), which exhibits a high propensity to self-assemble into a wide variety of fibril morphological polymorphs (47, 48). MUS:OT A NPs adsorbed selectively to the different tau peptide fibril polymorphic types – wide fibrils with a clear twist were densely decorated, while smooth and thin fibrils were not decorated by the NPs (Fig. 2F). This was observed even after prolonged periods of coincubation (Fig. S5) or with higher ratios of NPs to fibrils (Fig. 2G), with free and unbound NPs visible in the glassy ice, indicating that the different labeling patterns correlate with the different fibril surface properties rather than incubation conditions. Furthermore, within the labeled amyloid fraction, many polymorphs could be differentiated: straight and clear three-fold symmetric structures (Fig. 2H) and a whole range of two-fold twisted fibrils with different periodicity (Fig. 2I). These diverse fibril polymorphs could not be detected by negative staining or easily visualized by standard cryo-EM. As in the case of Aβ_40_, our NP-based procedure provided a rapid tool to establish the morphological diversity of the tau peptide fibrils.

To demonstrate that the NPs represent versatile cryo-EM markers for other amyloid-forming proteins, we incubated them with fibrils derived from different amyloidogenic proteins with different amino acid sequences that also form polymorphic fibrillar aggregates. To this end, we prepared fibrils from several proteins that have been linked to the pathogenesis of Alzheimer’s (tau) Parkinson’s (α-synuclein and C-terminally truncated α-synuclein (1-120)) and Huntington’s disease (mutant exon 1 of the huntingtin protein (residues 2 to 90), with 43 glutamines in the polyQ region – Httex1 43Q). Fig. 3 demonstrates that the NPs successfully label the fibrils produced from all these proteins, despite their sequence diversity and heterogeneous morphologies. Interestingly, compared to the Aβ_40_ and R2 fibrils, onto which the NPs bind to the fibril edges, both the tau and Httex1 43Q fibrils are decorated on the whole fibril surface.

**Fig. 3.**
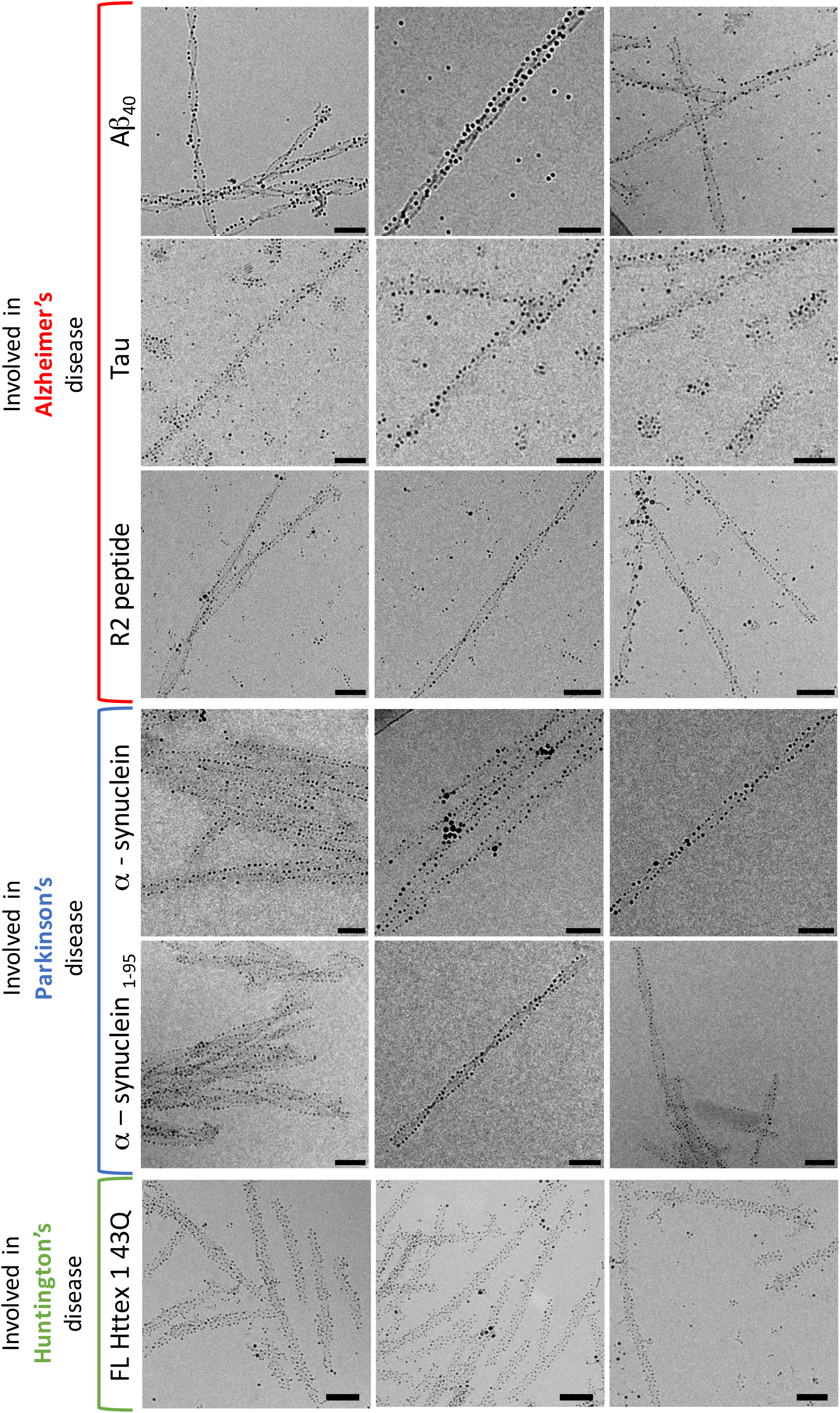
Examples of different amyloid fibrils decorated with MUS:OT NPs. Electron micrographs of fibrils prepared from proteins involved in Alzheimer’s disease (Aβ_40_, full-length tau and the tau-derived peptide R2), Parkinson’s disease (full-length and C-terminally truncated α-synuclein) and Huntington’s disease (mutant exon 1 of the Huntingtin protein, Httex1-43Q). All scale bars are 50 nm.

### MUS:OT NPs successfully label minute amounts of ex vivo samples and allow determination of polymorph distribution in human-derived samples

Finally, we sought to determine whether our NPs could be effective as a contrast agent in cryo-EM for human-derived samples. To achieve this goal, we used native tau PHFs isolated from the brain affected by AD and amyloid fibrils of an immunoglobulin λ light chain that were purified from the heart of a patient suffering from AL amyloidosis. We incubated both ex vivo fibril preparations with MUS:OT A NPs.

The cryo-EM images of AL amyloids show rather uniform and densely decorated fibrils with the crossovers clearly visible owing to the presence of the NPs, a feature that is not as evident in the normal cryo-EM and negative stain TEM images (Fig. 4A - C).

**Fig. 4.**
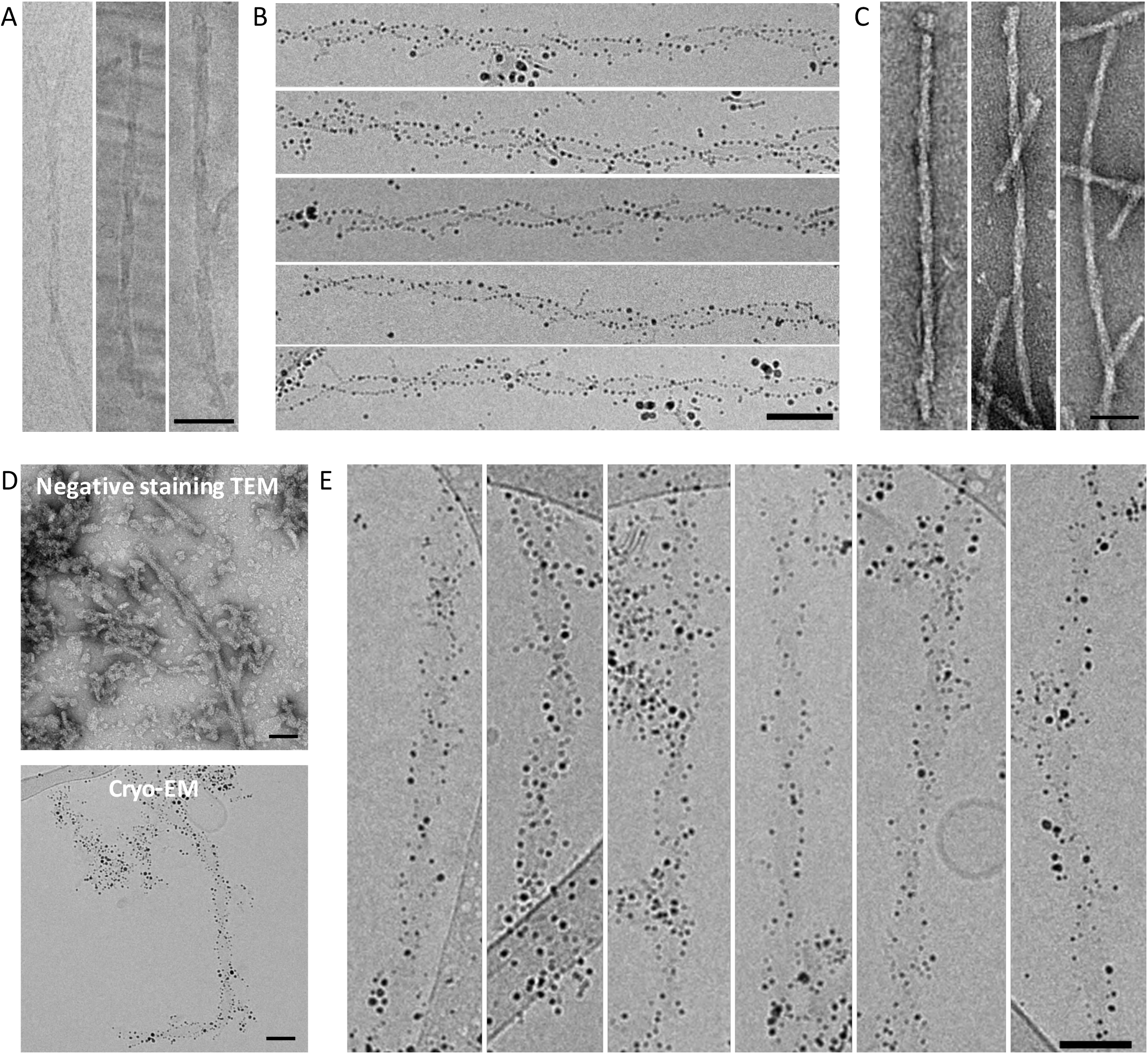
NP decoration of ex vivo PHFs and AL amyloid fibrils. A) Cryo-EM image of AL amyloid fibrils without NPs. B) A compilation of cryo-EM images of AL amyloid fibrils labeled with MUS:OT A NPs. C) AL fibrils imaged by negative stain TEM in the absence of NPs. D) Comparison between negative stain TEM images of PHFs and cryo-EM image of PHFs labeled with MUS:OT A NPs. E) A compilation of ex vivo PHF fibrils labeled with MUS:OT A NPs. Scale bars are 50 nm.

Similar data was obtained from enriched PHF samples from an AD postmortem brain. It is important to note that this sample still contained other nonamyloidogenic elements such as cell debris, proteins and other biological contaminants. The cryo-EM images indeed revealed the presence of many biological structures that were entangled with MUS:OT A gold NPs (Fig. S6). However, the PHFs were readily distinguishable from contaminants, which was due to the contrast provided by the NPs highlighting the characteristic fibrillar shape of a rather uniform double twisted morphology. (Fig. 4D – E).

For both native fibril samples, the NPs demonstrated specificity towards the edges of the amyloid fibrils and highlighted their delimitation, which allowed the observation of the morphological details and fibrillar characteristics. Furthermore, decoration with MUS:OT NPs allowed us to quantify specific features of the fibril polymorphs under hydrated conditions, such as the average periodicity width and length. We found that both ex vivo samples show a narrow periodicity range (Fig. 5B) when compared to the periodicity of the synthetic Aβ_40_ fibrils and to that of many other published fibrils prepared in vitro (49–51). The mean periodicity of the AL amyloids was equal to approximately 53.2 nm, with a standard deviation of 3.8 nm (N=73). The PHF fibrils derived from the brain of another patient had an average periodicity equal to 62.8 nm, with a standard deviation of 6.6 nm (N=18). This result contrasts with the values obtained for the in vitro obtained samples, which showed much higher standard deviations – 31.5 nm for Aβ_40_ fibrils (N=33) and 37.6 nm for tau peptide fibrils (N=20), with an average periodicity length of 76 nm and 95.8 nm, respectively. This contrast shows that the populations of fibrils present in each measured sample differ in morphology depending on the origin of the sample and demonstrates that human-derived amyloids are much more uniform than fibrils generated in vitro from recombinant or synthetic proteins and peptides. Further analysis of the periodicity showed that the length and width of the periodicity are positively correlated in the case of the fibrils produced in vitro. Amyloids derived from human tissue showed significant homogeneity of the periodicity width and no correlation between periodicity and fibril width (Fig. 5C).

**Fig. 5.**
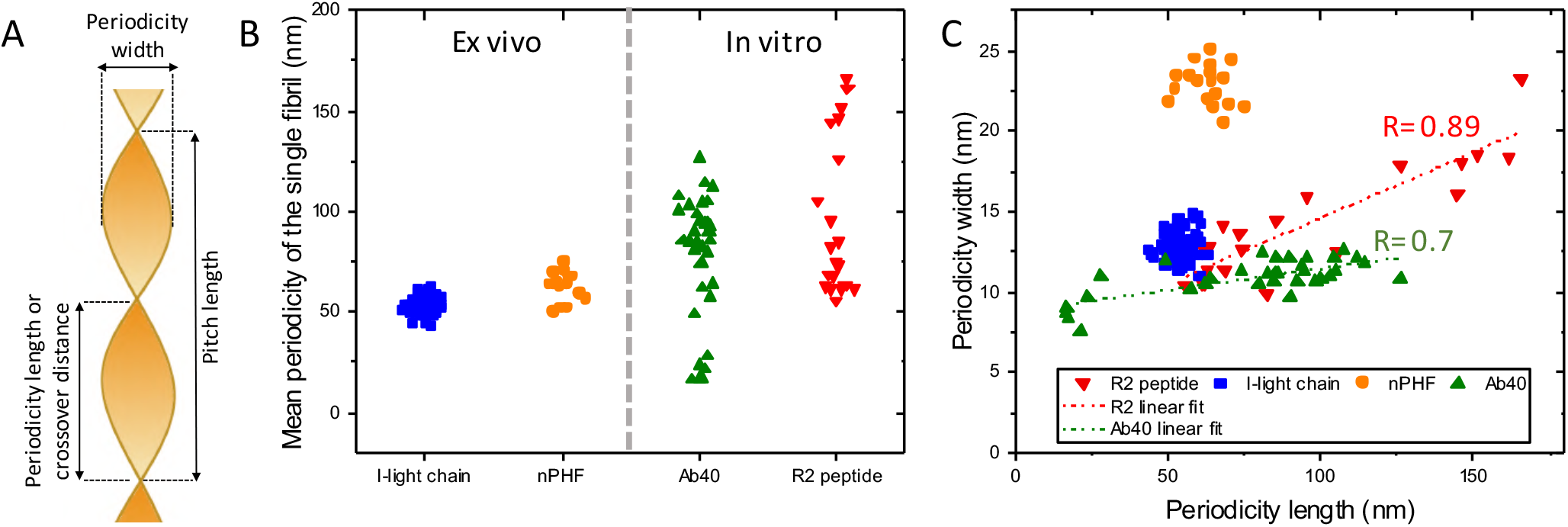
A) A schematic representation of the measured fibril characteristics – periodicity width, length or the crossover distance and pitch length. B) Distribution of the average crossover distance per fibril for the two types of ex vivo samples compared to the amyloids aggregated in vitro, highlighting the narrow distribution of crossover distance in the two ex vivo samples. C) Amyloid width plotted against crossover distance distribution. One dot represents the mean values obtained for the periodicity length and width of one fibril. The same figure enriched in the error bars can be found in the Supporting Information (Fig. S7).

Compared to existing methods, the coupling of MUS:OT NPs with cryo-EM allows for a straightforward examination of the entire population of fibrillar polymorphs in complex biological mixtures, requiring only a few microliters of sample. Moreover, the sample does not need to be extensively processed beforehand, as the fibrils bound to NPs are easily distinguished from other tissue components. Therefore, this method permits image analysis on minute amounts of sample of relatively low purity.

## Discussion

Increasing evidence suggests that amyloid fibril polymorphism underlines the emergence of various amyloid strains, which may explain the pathological diversity and possibly the clinical heterogeneity of amyloid diseases such as Alzheimer’s or Parkinson’s disease. For example, α-synuclein aggregates isolated from different types of synucleinopathies (e.g., Parkinson’s disease and multiple system atrophy) exhibited drastically different seeding capacity and induced different patterns of pathology spreading in in vitro neuronal cell models and animal models of synucleinopathies. Similar findings have been made for Aβ aggregates from AD brains and other amyloidogenic proteins. While these differences have been attributed to differences in the structural properties of the fibrils present in these samples, direct visualization and characterization of the aggregates has not been possible. This limitation is mainly due to the complexity of the brain homogenates and the minute content of fibrils and protein aggregates in these samples, which precludes the introduction of additional purification steps to allow their isolation and characterization. Although different approaches have been used to indirectly characterize such aggregates by taking advantage of their seeding capacity to try to amplify their structure using recombinant proteins as substrates, direct comparison of the amplified aggregates to the original seed has not been possible. Therefore, elucidation of the structural basis underlying the different properties associated with different fibril strains requires the development of new tools and experimental approaches that address these limitations and allow for direct visualization and structural characterization of fibrils in biological samples.

Recent advances in cryo-EM have enabled previously inaccessible insight into the structural properties of amyloid fibrils derived from recombinant and synthetic proteins of different sizes and even fibrils isolated from postmortem human tissues (11, 13, 52). One key step in the process of solving the structures of amyloids by cryo-EM involves the screening of sample conditions to identify conditions that reduce the structural and morphological diversity of the fibrils and/or favor the formation of specific populations of fibril structures. Therefore, the polymorphic diversity within a sample is not usually accounted for or reflected in the final structure. To address this limitation and allow for comprehensive and rapid analysis of fibril polymorphism, we propose the use of gold NPs as a potent tool to label and characterize fibril morphologies and polymorphism under hydrated conditions. Gold NPs have an electron-dense core that affords strong contrast under TEM and does not require any additional staining and/or processing to enhance the contrast. As such, we have developed an NP-based cryo-EM technique to simplify the analysis of fibril polymorphism derived from samples of different origin. The MUS:OT NPs exhibit a high propensity to attach to the edges of fibrils composed of various amyloid proteins, enabling rapid characterization of their polymorphic distribution in minute samples of increasing complexity. The improved contrast provided by the NP decoration also allows for quantitative image analysis, including the measurement of periodicity characteristics such as width and length. All of this can be done without extensive data collection and treatment. In a typical experiment, 3 microliters of fibril solution were sufficient to prepare a sample; hence, this technique allows for the labeling and characterization of fibrils in samples with low amounts of fibrils, e.g., purified fibrils isolated from postmortem tissues or biological fluids (e.g., plasma and cerebrospinal fluid, CSF).

The MUS:OT NPs appear to bind with different affinities to different fibril polymorphs. For example, in the case of the fibrils made of tau peptide fragments, we observed a different labeling pattern between twisted fibrils that were highly decorated and what appeared to be thin fibrils that were not decorated at all. These differences in labeling morphologically polymorphic species may indicate differences in surface properties and may be used in the future to separate specific polymorphs from a heterogeneous population using an NP-based purification strategy. Another mode of labeling was observed on the full-length recombinant tau, TDP 43 and Httex1 43Q fibrils, which were densely decorated along the whole surface, not only on the edges. This finding may be explained by the greater variability of exposed surfaces, which results in more adsorption sites for the NPs.

The analysis of recombinant and synthetic samples showed a wide range of structural polymorphs with variability in periodicity length and labeling efficacy. In Fig. 5C, we show an analysis that correlates the periodicity length and width for synthetic Aβ_40_ and recombinant tau peptides. The results are in agreement with the theory proposed by De los Rios (53), which assumes that the periodicity of the fibril depends on the number of protofilaments constituting the fibril. The correlation shown in Fig. 5C suggests a linear dependence of these two parameters for the amyloids aggregated in vitro. In contrast, the ex vivo samples analyzed here were composed of a rather homogenous fibril population, suggesting uniform protofibril number per mature amyloid. We observed a narrow distribution of the periodicity width and length in the AL and PHF amyloids, which showed remarkably conserved periodicity characteristics. This uniformness of the ex vivo fibrils contrasts with the amyloids derived from recombinant or synthetic proteins in cell-free systems, which revealed great polydispersity in terms of periodicity width and length. Our findings suggest that the mechanisms of amyloid formation in vitro may vary greatly from those occurring in vivo. An alternative explanation could be that amyloid formation in vivo is guided by complex interactions and conditions that are defined by the cellular or extracellular milieu (16) and are difficult to replicate in cell-free systems. The ex vivo fibril homogeneity could also be explained by the fact that each ex vivo sample was derived from a separate patient.

Previously published analysis of AL fibrils obtained from two patients showed that the fibrils formed two morphologically distinct aggregates that differed in width and periodicity length (30). Moreover, the two human-derived samples also presented distinct fibrillar characteristics, where one type of fibril revealed a width of approximately 11 nm and a pitch length distance of approximately 163 nm. The dimensions of these fibrils partially correspond to those of the AL fibrils that we analyzed. Our sample, however, revealed fibrils that were slightly wider and had a shorter pitch length. These differences may stem from the different methods used to obtain the micrographs, i.e., negative stain TEM and cryo-EM. It may also be caused by the fact that each patient develops fibrils of a different morphology.

Herein, we show that MUS:OT NPs efficiently label amyloid fibrils and allow for rapid profiling of their polymorphism. This labeling can be used in cryo-EM enabling direct detection and characterization of amyloid fibrils from ex vivo samples, with an image quality comparable to that achieved for amyloids obtained in vitro. Our findings also reveal striking morphological differences between the in vitro and ex vivo amyloid fibrils, consistent with emerging data pointing to the physiological milieu as the key determinant of amyloid fibril strains. Our method does not require elaborate enrichment, seeding, or any other form of extensive sample processing: the native fibril is directly observable in its solvated state. We believe that our NPs can be used as a complementary tool in studying the polymorphic distribution of amyloid fibrils by facilitating and accelerating sample screening, data acquisition and processing while requiring only minute amounts of sample. Moreover, the ability of the NPs to differently decorate polymorphic species open new doors in diagnostic strategies, as this would allow, for the first time, detailed characterization of fibril polymorphism of fibrils strains in biological samples (plasma and CSF) or in the identification of amyloids derived from postmortem brains or tissue biopsies.

## Materials and Methods

All compounds used in this study are detailed in the Materials and Methods section of the Supporting Information. Experimental procedures for ligand and NPs synthesis and characterization, amyloid fibrils preparation and incubation with NPs, microscopy measurement, circular dichroism and image analysis are described in Supporting Information.

## Supporting information

Supplementary Information complete

## Acknowledgments

The authors acknowledge AC Immune SA for the generous gift of native PHFs.

U.B.C. would like to acknowledge the funding received from the Swiss federal government agency (SERI) thanks to the EU H2020-MSCA-ITN project iSwitch (Grant Agreement No. 642196).

M.F. acknowledges support from the Deutsche Forschungsgemeinschaft (FA 456/27).

## References

1. Chiti F, Dobson CM (2017) Protein Misfolding, Amyloid Formation, and Human Disease: A Summary of Progress Over the Last Decade. Annu Rev Biochem 86:27–68.

2. Chiti F, Dobson CM (2006) Protein Misfolding, Functional Amyloid, and Human Disease. Annu Rev Biochem 75(1):333–366.

3. Knowles TPJ, Vendruscolo M, Dobson CM (2014) The amyloid state and its association with protein misfolding diseases. Nat Rev Mol Cell Biol 15(6):384–96.

4. Eisenberg D, Jucker M (2012) The Amyloid State of Proteins in Human Diseases. Cell 148(6):1188–1203.

5. Wechalekar AD, Gillmore JD, Hawkins PN (2016) Systemic amyloidosis. Lancet 387(10038):2641–2654.

6. Paravastu AK, Leapman RD, Yau W, Tycko R (2008) Molecular structural basis for polymorphism in Alzheimer’ s β-amyloid fibrils. Proc Natl Acad Sci U S A 105(47):18349–18354.

7. Kodali R, Williams AD, Chemuru S, Wetzel R (2010) Aβ(1-40) forms five distinct amyloid structures whose b-sheet contents and fibril stabilities are correlated. J Mol Biol 401(3):503–517.

8. Bousset L, et al. (2013) Structural and functional characterization of two alpha-synuclein strains. Nat Commun 4:2575.

9. Murzin AG, et al. (2019) Heparin-induced tau filaments are polymorphic and differ from those in Alzheimer’s and Pick’s diseases. Elife 8:1–24.

10. Ni X, McGlinchey RP, Jiang J, Lee JC (2019) Structural Insights into α-Synuclein Fibril Polymorphism: Effects of Parkinson’s Disease-Related C-Terminal Truncations. J Mol Biol 431(19):3913–3919.

11. Meinhardt J, Sachse C, Hortschansky P, Grigorieff N, Fändrich M (2009) Aβ(1-40) Fibril Polymorphism Implies Diverse Interaction Patterns in Amyloid Fibrils. J Mol Biol 386(3):869–877.

12. Goldsbury CS, Cooper GJS, Goldie KN, Mu SA (1997) Polymorphic Fibrillar Assembly of Human Amylin. 27:17–27.

13. Fitzpatrick AWP, et al. (2017) Cryo-EM structures of tau filaments from Alzheimer’s disease. Nature 547(7662):185–190.

14. Annamalai K, et al. (2016) Polymorphism of Amyloid Fibrils in Vivo. Angew Chemie - Int Ed 55(15):4822–4825.

15. Guo JL, et al. (2013) Distinct a -Synuclein Strains Differentially Promote Tau Inclusions in Neurons. Cell 154(1):103–117.

16. Peng C, et al. (2018) Cellular milieu imparts distinct pathological α-synuclein strains in α-synucleinopathies. Nature 557:558–563.

17. Peng C, Gathagan RJ, Lee VMY (2018) Distinct α-Synuclein strains and implications for heterogeneity among α-Synucleinopathies. Neurobiol Dis 109:209–218.

18. Peelaerts W, Bousset L, Baekelandt V, Melki R (2018) ɑ-Synuclein strains and seeding in Parkinson’s disease, incidental Lewy body disease, dementia with Lewy bodies and multiple system atrophy: similarities and differences. Cell Tissue Res 373(1):195–212.

19. Qiang W, Yau W-M, Lu J-X, Collinge J, Tycko R (2017) Structural variation in amyloid-β fibrils from Alzheimer’s disease clinical subtypes. Nature 541(7636):217–221.

20. Lu J-X, et al. (2013) Molecular Structure of β-Amyloid Fibrils in Alzheimer’s Disease Brain Tissue. Cell 154(6):1257–1268.

21. Sanders DW, et al. (2014) Distinct Tau Prion Strains Propagate in Cells and Mice and Define Different Tauopathies. Neuron 82(6):1271–1288.

22. Tycko R (2015) Amyloid Polymorphism: Structural Basis and Neurobiological Relevance. Neuron 86(3):632–645.

23. Adamcik J, Mezzenga R (2018) Amyloid Polymorphism in the Protein Folding and Aggregation Energy Landscape. Angew Chemie - Int Ed 57(28):8370–8382.

24. Eisenberg DS, Sawaya MR (2017) Structural Studies of Amyloid Proteins at the Molecular Level. Annu Rev Biochem 86(1):69–95.

25. Eichner T, Radford SE (2011) A Diversity of Assembly Mechanisms of a Generic Amyloid Fold. Mol Cell 43(1):8–18.

26. Zhu M, Souillac PO, Ionescu-Zanetti C, Carter SA, Fink AL (2002) Surface-catalyzed amyloid fibril formation. J Biol Chem 277(52):50914–50922.

27. Petkova AT, et al. (2005) Self-Propagating, Molecular-Level Polymorphism in Alzheimer’ s β-Amyloid Fibrils. Science (80-) 307:261–265.

28. Gath J, et al. (2014) Unlike twins: An NMR comparison of two α-synuclein polymorphs featuring different toxicity. PLoS One 9(3):1–11.

29. Fändrich M, Schmidt M, Grigorieff N (2011) Recent progress in understanding Alzheimer’s β-amyloid structures. Trends Biochem Sci 36(6):338–345.

30. Annamalai K, et al. (2016) Polymorphism of Amyloid Fibrils In Vivo. Angew Chemie - Int Ed (55):4822–4825.

31. Ohi M, Li Y, Cheng Y, Walz T (2004) Negative staining and image classification— powerful tools in modern electron microscopy. Biol Proced Online 6(1):23–34.

32. Vilar M, et al. (2008) The fold of alpha-synuclein fibrils. Proc Natl Acad Sci U S A 105(25):8637–42.

33. Page Faulk W, Malcolm Taylor G (1971) Communication to the editors. An immunocolloid method for the electron microscope. Immunochemistry 8(11):1081–1083.

34. Reig S, et al. (1995) Immunogold labelling of paired helical filaments and amyloid fibrils by specific monoclonal and polyclonal antibodies. Acta Neuropathol 90(5):441–447.

35. Conway KA, Harper JD, Lansbury PT (2000) Fibrils formed in vitro from α-synuclein and two mutant forms linked to Parkinson’s disease are typical amyloid. Biochemistry 39(10):2552–2563.

36. Hermann R, Walther P, Müller M (1996) Immunogold labeling in scanning electron microscopy. Histochem Cell Biol 106(1):31–39.

37. Skaat H, Margel S (2009) Synthesis of fluorescent-maghemite nanoparticles as multimodal imaging agents for amyloid-β fibrils detection and removal by a magnetic field. Biochem Biophys Res Commun 386(4):645–649.

38. Skaat H, Sorci M, Belfort G, Margel S (2009) Effect of maghemite nanoparticles on insulin amyloid fibril formation: Selective labeling, kinetics, and fibril removal by a magnetic field. J Biomed Mater Res - Part A 91(2):342–351.

39. Kumar J, et al. (2018) Detection of amyloid fibrils in Parkinson’s disease using plasmonic chirality. Proc Natl Acad Sci 115(13):3225–3230.

40. Liao YH, Chang YJ, Yoshiike Y, Chang YC, Chen YR (2012) Negatively charged gold nanoparticles inhibit Alzheimer’s amyloid-β fibrillization, induce fibril dissociation, and mitigate neurotoxicity. Small 8(23):3631–3639.

41. Yoshiike Y, Akagi T, Takashima A (2007) Surface structure of amyloid-β fibrils contributes to cytotoxicity. Biochemistry 46(34):9805–9812.

42. Zheng N, Fan J, Stucky GD (2006) One-step one-phase synthesis of monodisperse noble-metallic nanoparticles and their colloidal crystals. J Am Chem Soc 128(20):6550–6551.

43. Ong Q, Luo Z, Stellacci F (2017) Characterization of Ligand Shell for Mixed-Ligand Coated Gold Nanoparticles. Acc Chem Res 50(8):1911–1919.

44. Wilson AC, Dugger BN, Dickson DW, Wang D (2011) TDP-43 in aging and Alzheimer’s disease ‒a review. Int J Clin Exp Pathol 4(2):147–155.

45. Wang Y, Mandelkow E (2015) Tau in physiology and pathology. Nat Rev Neurosci 17(1):22–35.

46. Ganguly P, et al. (2015) Tau assembly: The dominant role of PHF6 (VQIVYK) in microtubule binding region repeat R3. J Phys Chem B 119(13):4582–4593.

47. Margittai M, Langen R (2006) Side chain-dependent stacking modulates tau filament structure. J Biol Chem 281(49):37820–37827.

48. Minoura K, et al. (2004) Different associational and conformational behaviors between the second and third repeat fragments in the tau microtubule-binding domain. Eur J Biochem 271(3):545–552.

49. Bedrood S, et al. (2012) Fibril structure of human islet amyloid polypeptide. J Biol Chem 287(8):5235–5241.

50. Adamcik J, et al. (2010) Understanding amyloid aggregation by statistical analysis of atomic force microscopy images. Nat Nanotechnol 5(6):423–428.

51. Watanabe-Nakayama T, et al. (2016) High-speed atomic force microscopy reveals structural dynamics of amyloid β _1–42_ aggregates. Proc Natl Acad Sci 113(21):5835–5840.

52. Jiménez JL, Tennent G, Pepys M, Saibil HR (2001) Structural diversity of ex vivo amyloid fibrils studied by cryo-electron microscopy. J Mol Biol 311(2):241–247.

53. Assenza S, Adamcik J, Mezzenga R, De Los Rios P (2014) Universal Behavior in the Mesoscale Properties of Amyloid Fibrils. Phys Rev Lett 113(26):1–5.

