## Supplementary Information complete for "Unraveling the Complexity of Amyloid Polymorphism Using Gold Nanoparticles and Cryo-EM"

PNAS
www.pnas.org

Supplementary Information for

**Deciphering Amyloid Polymorphism Using Gold Nanoparticles**

Urszula Cendrowska, Paulo Jacob Silva, Nadine Ait-Bouziad, Marie Müller, Zekiye Pelin Guven, Sophie Vieweg, Anass Chiki, Lynn Radamaker, Senthil Thangaraj, Marcus Fändrich, Francesco Tavanti, Maria Cristina Menziani, Alfredo Alexander-Katz, Francesco Stellacci, Hilal A. Lashuel

Hilal A. Lashuel

Francesco Stellacci


**This PDF file includes:**

Supplementary text
Figures S1 to S8
References for SI reference citation

### Supplementary Information text

### Supplementary Information - Materials and Methods

Unless otherwise indicated, all chemicals were purchased from Sigma-Aldrich. All $^1$H-NMR spectra were acquired on a Bruker Avance 400 MHz or Avance III 400 MHz spectrometer.

#### Ligands

##### 11-Mercapto-1-undecanesulfonate (MUS) ligand synthesis

The MUS ligand was synthesized in three steps. First, sodium undec-10-enesulfonate was synthesized; then, sodium 11-acetylthio-undecanesulfonate and finally 11-mercapto-1-undecanesulfonate (MUS) were synthesized.

1. *Sodium undec-10-enesulfonate synthesis*

11-Bromo-1-undecene (25 ml, 111.975 mmol), sodium sulfite ($Na_2SO_3$, 28.75 g, 227.92 mmol) and benzyltriethylammonium bromide (10 mg) were added to a mixture of 200 ml of methanol and 450 ml of DI water (4:9 MeOH:H2O ratio) in a 1 l round bottom flask. The mixture was refluxed at 102°C for 48 h. The mixture was extracted with diethyl ether 5 times (5 x 400 ml), and the aqueous phase was evaporated in a rotary evaporator. The white powder was dried under high vacuum, suspended in pure methanol and filtered. The methanolic solution was evaporated, and the process was repeated twice to decrease the amount of inorganic salts present in the sample. $^1$H-NMR ($D_2O$): 5.76 (m, 1H), 4.78 (m, 2H), 2.69 (t, 2H), 1.53 (m, 2H), 1.11 (br s, 12H).

2. *Sodium 11-acetylthio-undecanesulfonate synthesis*

Sodium undec-10-enesulfonate (33 g, 147.807 mmol) was dissolved in 500 ml of methanol. A 2.6-fold excess of thioacetic acid (27.324 ml, 384.3 mmol) was added to the solution, and the mixture was stirred under a UV lamp overnight (12 h). The solution was evaporated in a rotary evaporator until the solid residue turned orange-red. The solid was washed with diethyl ether until no colored material could be removed. The solid was dried under high vacuum and then dissolved in methanol to produce a yellow solution. Next, 3 g of carbon black was added to the solution, which was vigorously mixed, and the mixture was filtered through Celite in a fluted filter paper. The solvent of the clear filtrate was completely evaporated, and white solid was collected. $^1$H-NMR ($D_2O$): 2.69 (t, 4H), 2.17 (s, 3H), 1.53 (m, 2H), 1.39 (m, 2H), 1.11 (br s, 14H).

3. *11-Mercapto-1-undecanesulfonate (MUS) synthesis*

Sodium 11-acetylthio-undecanesulfonate was refluxed at 102°C in 400 ml of 1 M hydrochloric acid (HCl) for 12 h. Then, 200 ml of 1 M sodium hydroxide (NaOH) was added to the final solution, and an additional 400 ml of DI water was added to create a 1 l volume. The clear solution was kept at 4°C and crystallized overnight. The viscous white product was centrifuged down in 50 ml Falcon tubes and dried under high vacuum. An approximately 30% yield of MeOH-soluble MUS was collected from this purification step. More material was extracted from the supernatant of the centrifugation step by reducing the volume and keeping it at 4°C. $^1$H-NMR ($D_2O$): 2.69 (t, 4H), 2.34 (t, 3H), 1.53 (m, 2H), 1.39 (m, 2H), 1.11 (br s, 14H). Calculated mass 290.42 g/mol.

**N,N,N-Trimethyl(11-mercaptoundecyl)ammonium chloride (TMA) ligand synthesis**

This ligand was synthesized in two steps. First, N,N,N-trimethyl-10-undecenylammonium chloride was synthesized, and then, N,N,N-trimethyl(11-mercaptoundecyl)ammonium chloride was synthesized.

1. *N,N,N-Trimethyl-10-undecenylammonium chloride synthesis*

11-Bromo-1-undecene (25 ml, 111.975 mmol) was added to 300 ml of an ethanolic trimethylamine solution (31-35 wt. % in ethanol, 4.2 M) and stirred for 2 days at room temperature. The solvent was evaporated, and the yellow powder was dissolved in ~50 ml of methylene chloride (DCM) and precipitated into hexane (500 ml) in a 1 L beaker, yielding 30 g of white solid.

2. *N,N,N-Trimethyl(11-mercaptoundecyl)ammonium chloride*

N,N,N-Trimethyl(10-undecenyl)ammonium bromide (30 g, 102.6 mmol) and thioacetic acid (21 ml, 308 mmol, ACROS) in 500 ml were mixed and stirred under a UV lamp overnight (12 h). The volatiles were removed using a rotary evaporator until the product became orange-red. The residue was washed several times with diethyl ether until no more orange byproduct could be removed. The product was dried under high vacuum and then dissolved in 300 ml of methanol, to which ~3 g of carbon black was added, and the mixture was vigorously mixed, followed by filtration through Celite in a fluted filter paper. The clear solution was evaporated, yielding 32 g of white powder. This product was then dissolved in 400 ml of 1 M HCl and refluxed at 102°C overnight (~12 h). The pH was increased by the addition of 100 ml of 1 M NaOH, followed by the addition of 400 ml of MilliQ water, and the solution was placed inside a refrigerator at 4°C. Thin elongated crystals grew and were collected via centrifugation-assisted decantation. After drying, ~10 g of a "shiny" crystalline powder was collected. 1H-NMR ($D_2O$): 1.24-1.49 (m, 14H), 1.53- 1.66 (m, 2H), 1.67-1.84 (m, 2H), 2.52 (t, 8 Hz, 2H), 3.12 (s, 9H), 3.30-3.41 (m, 2H).

**3-[(11-Mercapto-undecyl)-N,N-dimethylamino]propane-1-sulfonate (ZW) synthesis**

11-Bromo-1-undecene (15 ml, 68.4 mmol) was added to a solution of 2 M dimethylamine in oxolane (THF) (AcroSeal™, ACROS Organics™), and the reaction mixture was stirred for 48 h at room temperature. The volatiles were evaporated in a rotary evaporator. A yellow oil remained, to which 200 ml of 1 M NaOH was added, followed by extraction with DCM. The DCM phase was separated, dried with anhydrous sodium sulfate, filtered, and concentrated in vacuo as a yellow oil. Next, 6.02 g of this oil was added to 100 ml of dry acetone (AcroSeal™, ACROS Organics™), followed by the addition of 1,3-propanesultone (1.6 ml, 38.25 mmol): the reaction mixture was stirred at room temperature for two days. The white precipitate was filtered, and the resulting solid was washed with excess acetone and then dried under high vacuum. Then, 4 g (~11.68 mmol) of the powder was dissolved in methanol with thioacetic acid (1.426 ml, 20 mmol) and stirred at room temperature under a UV lamp. The volatiles were removed in a rotary evaporator, yielding a yellow oil. Next, 20 ml of methanol was added to this oil, and this solution was added dropwise into 400 ml of dry acetone (AcroSeal™, ACROS Organics™). The resulting white precipitate was filtered using vacuum filtration and dried under vacuum. A solution of methanolic HCl was prepared by adding 3.56 ml (50 mmol) of acetyl chloride to 50 ml of methanol. The powder was added to this mixture and refluxed overnight. The volatiles were evaporated in a rotary evaporator, which produced a yellow oil. Toluene was added and evaporated from this oil until a pale yellow powder formed (~3 g).

1-Octanethiol (OT) and 11-mercaptoundecylphosphoric acid (MUP) were purchased from Sigma-Aldrich.

### Nanoparticle synthesis and characterization

MUS, MUP and MUS:OT NPs were synthesized according to the one-phase method(1) with modifications in the particle purification step. TMA and ZW particles were synthesized according to a modified Stucky synthesis (2). All glassware was cleaned with fresh aqua regia (HCl:HNO3 3:1) before synthesis.

#### One-phase MUS and MUS:OT NP synthesis

In a 500 ml round-bottom flask, 118.15 mg (0.3 mmol) of gold salt ($HAuCl_4 \cdot 3H_2O$) in 200 ml of ethanol was stirred until the gold salt was completely dissolved. In a separate glass vial, MUS or the mixture of MUS:OT was dissolved in 15 ml of methanol, aided by sonication. Usually, to reach a 1:1 feed ratio of the ligands, we added 65.5 mg of MUS and 39 μl of OT. The desired thiol ligand mixture was then added to the gold salt dissolved in ethanol while stirring. The solution of gold salt and thiolated ligands was stirred together for approximately 15 min. During that time, the color of the solution changed from translucent to turbid yellow, indicating the formation of gold–thiolate complexes. Then, a filtered, saturated solution of 500 mg of sodium borohydride ($NaBH_4$) dissolved in 100 ml of ethanol was added dropwise. The addition of $NaBH_4$ was adjusted such that the whole process took 1 h to avoid quick reduction. The solution was stirred for another hour. After that time, the reaction was assumed to be finished, and the vessel was closed with the septum pierced with a needle to allow controlled $H_2$ gas release. The flask was then placed in the refrigerator and kept at 4˚C overnight. The next day, a black precipitate was collected via decantation. This residue was washed several times (5 times each step) with ethanol and dried under vacuum to remove ethanol. To completely remove unbound species, particles were centrifuged several times with MilliQ water using Amicon® Ultra-15 centrifugal filter devices (10k or 30k NMWL). The particles were then suspended in a small amount of water (~2 ml) and freeze-dried.

**One-phase MUP NP synthesis**

Separately, 12-mercaptoundecylphosphoric acid (MUP) (255 mg, 0.9 mmol) and $NaBH_4$ (2 g) were dissolved in ethanol (20 ml and 200 ml, respectively, Fluka, Puriss > 99.8%). Both solutions were then sonicated to aid dissolution and filtered to remove any insoluble residue. In a third container, gold(III) chloride trihydrate (354 mg, 0.9 mmol) was dissolved in ethanol (200 ml, Fluka). The MUP ligand solution was then added to the gold salt solution with stirring for 10 min. The $NaBH_4$ solution was then added dropwise to this vigorously stirred solution. After complete addition of the reducing agent, the mixture was stirred for 1 h, and then, the reaction flask was stored overnight at 4°C to precipitate the NPs. The NPs were then spun down (5000 rpm). The supernatant was removed, and the NPs were redispersed in 45 ml of ethanol. Ethanol washing was repeated 3 times. The residue was then dispersed in water (15 ml) and filtered through Amicon® Ultra-50 centrifugal filter devices (30k MW cutoff) to further wash the particles. This step was repeated until the water removed no longer foamed when shaken. The NP solution was then dialyzed (8k MW cutoff) against water for 2 weeks with water changes once per day. At the end of this process, the particles appeared less soluble in water, so they were further dialyzed (tubing of 8k MW cutoff) against aqueous NaOH (pH 12) for two days before dialysis against pure water for 1 day. The particles were then freeze-dried to yield a purple powder.

**Stucky synthesis of TMA and ZW NPs**

A 1:1 mixture of ethanol and toluene was prepared in a 250 ml round-bottom flask. Three different 20 ml aliquots of this mixture were used to completely dissolve the reagents in separate vials: (i) 277.7 mg (0.56 mmol) of chloro (triphenylphosphine) gold(I), (ii) 1.2 mmol of TMA or ZW ligand and (iii) 142.3 mg (1.5 mmol) of borane tert-butylamine complex. Dissolution was completed by sonication for 15 min at room temperature. The gold-salt solution was added to a round-bottom flask, followed by the addition of the ligand solution. The mixture was stirred at 800 rpm for 10 min at room temperature. The reducing agent solution was then added, and the flask was connected to a condenser in an oil bath at 125°C (800 rpm) for 1.5 h. Afterwards, the reaction flask was cooled while stirring (800 rpm). Here, the methodology for ZW and TMA nanoparticle synthesis departed: (i) The ZW NPs precipitated like the MUS and MUS:OT particles, and the cleaning procedure followed exactly that described above. (ii) The TMA NPs did not precipitate; they remained stably soluble in the mixture. Thus, the volume was decreased in a rotary evaporator, which decreased the ethanol content of the mixture; once toluene was in a large enough excess, the NPs precipitated. Since the TMA NPs are soluble in ethanol, they were washed by centrifugation using diethyl ether and toluene. The pellet was dried and dissolved in MilliQ water, followed by Amicon® Ultra-15 centrifuge-assisted dialysis (10k or 30k MW cutoff). The particles were then suspended in a small amount of water (~2 ml) and freeze-dried.

### Nanoparticle characterization

The NPs were systematically characterized using TEM, $^{1}$H-NMR spectroscopy and thermogravimetric analysis (TGA), as presented in Fig. S1 (MUS:OT A as an example). TEM allowed for the quantification of the NP size distribution. Nuclear magnetic resonance spectroscopy was used both to assess the presence of unbound ligands and to determine the ligand-shell composition. TGA was performed to analyze the ligand density on the NPs.

**Cleanliness of the NPs**

To control the level of impurities in the sample, $^{1}$H-NMR analysis was performed. Five milligrams of the NP powder was dissolved in 600 µl of $D_2O$. The absence of sharp peaks in the NMR spectrum indicates the absence of impurities in the solution.

**Ligand-shell composition determination**

The ratio between protective ligands in the case of the mixed ligand nanoparticles, such as MUS:OT, was assessed with the use of $^{1}$H-NMR spectroscopy. An etching solution of 15 mg of iodine (Acros) in 100 ml of MeOD-d4 (Sigma) was prepared. Between 1 and 5 mg of NPs were suspended in 0.6 ml of the etchant mixture for 30 min under sonication. After the NMR spectrum for the etched solution was obtained, the ligand ratio was calculated according to the integrals of the given peaks.

**Thermogravimetric analysis (TGA)**

TGA offers an estimate of the organic content of the sample. It can also be used to estimate the relative abundance of the different ligands in our NPs, as they have discernible thermal decompositions. The equipment used was a TGA 4000 system from Perkin Elmer. Between 2 and 8 mg of the NP powder was placed into a TGA crucible. The temperature was increased to 900°C with heating at 5°C per min. The organic ligands degrade in time as the temperature increases, leaving the gold core of the NPs. Degradation of the ligands causes weight loss, which is monitored and plotted against the temperature, enabling the monitoring of ligand desorption with time. The difference between the beginning and the final weight for the sample allows for the determination of the ligand density on the NPs.

**Representative TEM analysis of NPs**

A drop of 4 µl of NPs (0.1 – 0.5 mg/ml) was deposited onto a 400-mesh carbon-supported copper grid and left to dry. All TEM images were acquired using an FEI TALOS™ electron microscope with an acceleration voltage of 200 kV and equipped with a Ceta CCD camera.

Images of the NPs were analyzed using Fiji software, and their diameter was calculated using a homemade script compatible with this software.

### Amyloid fibril preparation

#### Specific amyloid fibril preparations

*$A\beta_{40}$ fibrils*

$A\beta_{40}$ samples were prepared according to the previously reported protocol (3). $A\beta_{40}$ was purchased from ChinaPeptides. The lyophilized material was dissolved in a 1:1 mixture of 0.1% $NH_4OH$ and 100 mM Tris buffer (with 0.02% $NaN_3$ and at pH 7.4) at approximately 1 mg/ml. The solutions were ultracentrifuged for 1 h at 366941 RCF (g) at 4°C in a Beckman ultracentrifuge. The upper half of the supernatant was collected, and the peptide concentration was determined using its extinction coefficient at 280 nm (1490 $M^{-1}cm^{-1}$). A Perkin Elmer UV-vis or a Tecan plate reader was used to perform these measurements. The supernatant was then diluted to the desired concentration (usually between 5 and 50 µM) with the PBS (pH = 7.4) described above. Fibrils were grown either in a Tecan plate reader or inside 1.5 ml Eppendorf tubes in a thermomixer (Eppendorf). Fibrils were grown under quiescent conditions, except agitated fibrils, which were shaken at 600 rpm.

*Wild-type $\alpha$-synuclein*

Human full-length $\alpha$-synuclein was purchased from rPeptides and used as received. To 1 mg of lyophilized powder, 1 ml of TBS buffer was added, and the solution was filtered through a 100 kDa MW cutoff Microcon filter (13000 g, 15 min, 4°C). The concentration was adjusted to 40 µM using the extinction coefficient at 275 nm (5974 $M^{-1}cm^{-1}$). Fibrils were grown over the period of one week at 1000 rpm and 37°C.

*Modified $\alpha$-synuclein*

Human truncated $\alpha$-synuclein (1-120) in pT7-7 was expressed in *E. coli* strain BL21, purified and characterized as described by Fauvet et al. (4) except that the anion exchange chromatography step was replaced by a cation exchange chromatography step due to the lack of the negatively charged C-terminal domain. The fibrils were prepared as described for the wild type protein.

*Wild-type tau*

Human full-length tau (isoform 4R2N, 441 amino acid) in pET-15b was expressed in *E. coli* strain BL21. K19 and the purification was adapted from ref (5). Briefly, cells were pelleted and broken by sonication in lysis buffer (3 M urea in 10 mM MES, pH 6.5, 1 mM DTT, 1 mM EDTA, 1 mM PMSF). After centrifugation at 150,000 × g for 1 h at 4°C, 1% (w/v) streptomycin sulfate was added to the supernatant, and the solution was stirred for 90 min at 4°C. After centrifugation at 27,000 × g for 1 h at 4°C, the supernatant was dialyzed overnight at 4°C in ion exchange (IEX) buffer A (10 mM MES, pH 6.5, 20 mM NaCl, 1 mM DTT, 1 mM EDTA). The supernatant was filtered and loaded on a cation exchange column (MonoS, GE Healthcare), and the protein was eluted using a salt gradient (increasing the NaCl concentration of IEX buffer A from 20 mM to 1 M NaCl over 20 column volumes). Fractions containing the proteins were dialyzed overnight against acetic acid buffer (5% acetic acid in water) and loaded on a reversed-phase HPLC C4 column (PROTO 300 C4 10 µm, Higgins Analytical; buffer A: 0.1% TFA in water, buffer B: 0.1% TFA in acetonitrile), and the protein was eluted using a gradient from 30 to 40% buffer B over 40 min (15 ml/min).

A solution of 20 µM of the longest isoform of human tau (tau441) was prepared by adding 1.7 ml of PBS (pH 7.4) to 1.6 mg of the lyophilized powder. To the largest 0.9 ml aliquot, heparin was added to a final concentration of 4 µM, aiming for a tau-to-heparin 4:1 ratio. The solution was shaken for 48 h until mature fibrils were obtained.

*R2 peptide*

The tau-derived R2 peptide 275 – VQIINKKLDLSNVQSKCGSKDNIKHV – 300, (numbering according to human tau isoform 2) was purchased from CisBio. R2 amyloid fibrils were prepared by incubating 100 µM of the R2 peptide in the presence of 1:4 (mol:mol) of heparin in 10 mM phosphate buffer pH 7.4, 50 mM NaF and 0.5 mM freshly dissolved DTT at 37°C under quiescent conditions for at least 12 h.

*Huntingtin exon 1 (Httex 1)*

Both fibril types – N-truncated 43Q (with the first 17 amino acids removed) and full-length 43Q Httex1 – were prepared according to a previously reported protocol (6). Briefly, Httex1 with polyQ repeats equal to 43Q was synthesized using an intein-based strategy that allows for the production of native tag-free huntingtin exon 1. Fibrils were formed by keeping the solution of monomers at 37°C without shaking.

*AL amyloid fibrils*

Heart tissue was collected at the University Clinic Heidelberg, Germany, from a 51-year-old male patient who had to undergo a heart transplantation due to systemic AL amyloidosis with severe cardiac involvement. The method for fibril extraction from human tissue was based on a previously established protocol(7), making use of the water solubility of fibrils, with small amendments. A total of 250 mg of frozen heart tissue was kept on ice shortly and then diced into fine pieces using a scalpel. Next, 0.5 ml of trisaminomethane (Tris) calcium buffer (20 mM Tris, 138 mM NaCl, 2 mM $CaCl_2$, 0.1% $NaN_3$, storing condition 4°C, pH 8.0) was added to the diced tissue material. The sample was mixed using a Kontes pellet pestle for 10 s in a pulsating manner (1 s on, 1 s off) and centrifuged (6000 rpm, 5 min, 4°C). The supernatant was stored. This cycle of resuspension, mixing, centrifuging and storing was repeated 5 times, and a clear supernatant was obtained. After the last centrifugation step, the pellet was resuspended in 1 ml of freshly prepared 5 mg ml$^{-1}$ *Clostridium histolyticum* collagenase (Sigma) and ethylenediaminetetraacetic acid (EDTA)-free protease inhibitor (Roche) (1 tablet in 7 ml of Tris calcium buffer) in Tris calcium buffer and incubated for 17 h at 37°C under constant agitation (1100 /min, horizontal position of the tube). Afterwards, the sample was centrifuged (6000 rpm, 20°C, 30 min). The supernatant was stored, and the pellet was washed 10 times by the addition of 0.5 ml of ice-cold Tris EDTA buffer (20 mM Tris, 140 mM NaCl, 10 mM EDTA, 0.1% $NaN_3$, storing condition 4°C, pH 8.0), followed by mixing with the Kontes pellet pestle and centrifugation (6000 rpm, 5 min, 4°C). For each step, the supernatant was stored. The same washing procedure was repeated with MilliQ water instead of Tris EDTA buffer for 10 steps. The collection of human material was conducted under the approval of the ethical committee at Heidelberg University, while the extraction of fibrils from the tissue was approved by the ethical committee of Ulm University.

**Incubation of the amyloid fibrils with NPs**

Recombinant amyloid fibrils prepared as described above were dispersed in their corresponding buffer and incubated at 37°C in a thermomixer with a water solution of NPs. The final concentration of the NPs ranged from 0.1 mg/ml to 0.3 mg/ml. The labeling speed was assessed by cryo-EM and depended on the amyloid type (e.g., amyloids made from the R2 protein become decorated much faster than $A\beta_{40}$), but 24-48 h of incubation with the NPs was sufficient for most of the fibrils we worked with, and all types of amyloid fibrils were incubated with NPs for 24 h unless stated otherwise in the paper. Shaking of the sample enabled continuous movement of all of the elements in the sample and gave momentum to the NPs, which greatly improved the decoration process. We found that the optimal shaking speed was in the range of 300 rpm – 600 rpm. Most of the presented amyloids were shaken with NPs at 300 rpm except $A\beta_{40}$, which usually requires more vigorous shaking to obtain sufficient decoration and was incubated with NPs at 600 rpm shaking. A rotation slower than 300 rpm was not efficient, and a higher speed risked breaking the fibrils.

To limit any unwanted biological activity (e.g., from enzymes), the ex vivo samples were incubated at low temperature (20°C overnight) with a final concentration of NPs ranging from 0.1 mg/ml to 0.5 mg/ml. The nature of the sample (i.e., still containing other proteins and biological structures) required a higher concentration of NPs. During the incubation, agitation was applied (300 rpm). Incubation was performed for 24 h-48 h until sufficient coverage of the fibrils with NPs was obtained.

In the case of the PHFs, a dark sediment was observed at the bottom of the Eppendorf tube, suggesting strong adsorption of our nanomaterial to structures present in the sample. The sample was gently resuspended using a pipette to generate some agitation immediately prior to cryo grid preparation.

**Negative stain TEM of amyloids**

Recombinant samples were deposited onto glow-discharged 400-mesh carbon-supported copper grids for 1.5 min at room temperature. The grids were then blotted with filter paper, washed once with MilliQ water and stained with a 1% w/v uranyl acetate solution for 30 s. The blotted grids were air-dried and imaged using an FEI TALOS electron microscope at an acceleration voltage of 200 kV and equipped with a Ceta CCD camera. In the case of the ex vivo samples, 3.5 µl of each sample was applied onto glow-discharged Formvar/carbon-coated 200-mesh copper grids (Electron Microscopy Sciences) for 1 min. The grids were blotted with filter paper, washed twice with ultrapure water, washed once with uranyl formate 0.7% (w/V), stained with uranyl formate for 30 s, blotted and dried. The prepared specimens were inspected with a Tecnai Spirit BioTWIN system operated at 80 kV and equipped with an Eagle CCD camera.

**Cryo electron microscopy of amyloid and nanoparticle samples**

For cryo-EM microscopy, a droplet of amyloid fibrils suspended in buffer was deposited onto a Quantifoil® holey carbon grid or lacey carbon film (Electron Microscopy Sciences) and blotted to a thin (100-300 nm) layer of liquid that was flash frozen in liquid ethane using an FEI Vitrobot Mark IV. Imaging was performed using a Gatan single tilt cryo holder operated on an FEI Tecnai Spirit BioTWIN 80 kV transmission electron microscope in LowDose Mode to visualize the samples at an average exposure of 1-3 electrons/$^2$ on a Ceta camera.

**Circular dichroism**

Samples were analyzed on a J-815 CD spectrometer from Jasco using a 1-mm quartz cuvette. The CD spectra were acquired from 190–250 nm at room temperature, and 3-5 spectra of each sample were obtained and averaged. The obtained spectra were further processed by smoothing using a binomial filter with an iteration equal to 5. Data were acquired with data pitch 1 nm and bandwidth equal to 4 nm at a scanning speed of 200 nm/min.

CD spectra of mature $A\beta_{40}$ amyloid fibrils (20 µM) were obtained by incubating mature amyloid fibrils with NPs at a final concentration of 0.1 mg/ml, except for the control sample, where an equivalent volume of water was added instead of the NP solution. Mixtures were incubated at 37°C with shaking (600 rpm), and CD measurements were performed at the beginning of the incubation (0 h) and after 1, 16, 24 and 48 h of incubation.

CD spectra of monomeric $A\beta_{40}$ incubated with NPs were obtained using the same procedure. Monomeric $A\beta_{40}$ (35 µM) was incubated at 37°C with NPs at a final concentration of 0.1 mg/ml, except for the control sample, where an equivalent volume of water was added instead of the NP solution. The samples were agitated at 600 rpm. CD spectra were obtained at the beginning of the incubation (0 h) and after 1, 16, 24 and 48 h of incubation.

### Fibril crossover distance and width analysis

Micrographs of the amyloids decorated with NPs were analyzed using ImageJ software, where the crossover distance of the fibril was measured manually. For each fibril, the average crossover distance and width were calculated and plotted in the graphs using OriginPro software. In the case of the human-derived samples, 73 AL amyloid fibrils were evaluated from 40 acquired images, and 18 nPHF fibrils were measured on 18 micrographs. Recombinant sample data were obtained from 20 R2 fibrils spotted on 16 micrographs and 33 $A\beta_{40}$ amyloids visible on 26 images.

### Supplementary information – Results

#### Ligand screening

In order to screen for the best amyloid fibrils' labelling efficiency, we tested several different ligands that were composing the shell of our NPs. We found that zwitterionic ligands did not interact with fibrils, and cationic ligands (TMA) exhibited non-specific interactions that led to the formation of large fibril-NP aggregates (Fig. S2a.). NPs coated with negatively charged MUP ligands showed a tendency to aggregate in the solution, while NPs labeled with negatively charged MUS ligands successfully attached to the $A\beta_{40}$ amyloids, albeit in a sporadic and random manner (Fig. S2a). To improve the decoration obtained with these NPs, we modified the ligand shell composition by adding hydrophobic ligand OT to maximize hydrophobic contacts with the $A\beta_{40}$ fibrils. We found that a ratio of 4 to 1 between the negatively charged and hydrophobic ligands on the gold core (MUS:OT A) greatly improved the labeling of the amyloid fibrils under the negative staining TEM (Fig. S2b). Therefore, we established, that the mixture of OT and MUS ligands on the gold surface was the most efficient in labeling $A\beta_{40}$ fibrils and this ligand composition was applied in the further experiments.

#### Molecular dynamics

To investigate the nature of the MUS:OT NP interaction with $A\beta_{40}$ fibrils, we performed molecular dynamics simulations of a NP with a core diameter of 2 nm with both 100% MUS ligands and an MUS:OT ligand ratio of 7:3. The coated NPs are built according to previous work (8), while the $A\beta_{40}$ fibril structure was retrieved from the work of Petkova et al. (9), which is characteristic of the two-fold $A\beta_{40}$. The single $A\beta_{40}$ protofibril, which is a precursor aggregate of mature amyloid, was placed in a cubic box with the same length as that of the fibril to obtain a continuous, indefinitely long fibril, as previously done by Buchete et al. (10). A single NP was randomly placed in the simulation box so that the fibril and the NP were not in contact with each other. The spontaneous binding of monolayer-coated NPs onto the fibril was observed during the 100 ns-long simulations.

The NPs interact with the beta-1 region of the fibril ("β-1 binding"), establishing stable contacts with the hydrophobic amino acids from H14 to F20, as shown in Fig. S8. During the simulations involving the 2 nm 7:3 NP, we observed that the unordered tails in the $A\beta_{40}$ N-terminal region interact with the NP before binding to the fibril. In this case, up to 4 tails establish contacts with the NP, especially through A2 and F4 amino acids.

The binding site of both MUS NP and 7:3 NPs on the fibrils involves the stretch of amino acids from H14 to F20 ($^{14}$HQKLVFFA$^{21}$), in agreement with the predicted binding site ($^{16}$KLVFFA$^{21}$) for drugs and peptides reported in the literature (11, 12). This binding site is characterized by the presence of hydrophobic amino acids, suggesting that hydrophobic interactions play a relevant role in the binding of NPs.

A higher number of contacts, approximately 220, between the 7:3 NP and the fibril is observed due to the interaction with the hydrophobic amino acids on the fibril's tails with respect to the MUS NP, which has approximately 180 contacts. The estimation of the hydrophobic component of the free energy ($\Delta G_{phobic}$) shows that the value for the 7:3 NP (-197 ± 56 kcal/mol) is slightly higher than that for the MUS NP (-179 ± 47 kcal/mol) due to the hydrophobic contacts between the NP and $A\beta_{40}$.

Moreover, the MD simulations show that the NPs do not change the secondary structure of the $A\beta_{40}$ fibrils during the simulation time, as shown by the CD analysis.

Supplementary Information – Figures and Legends

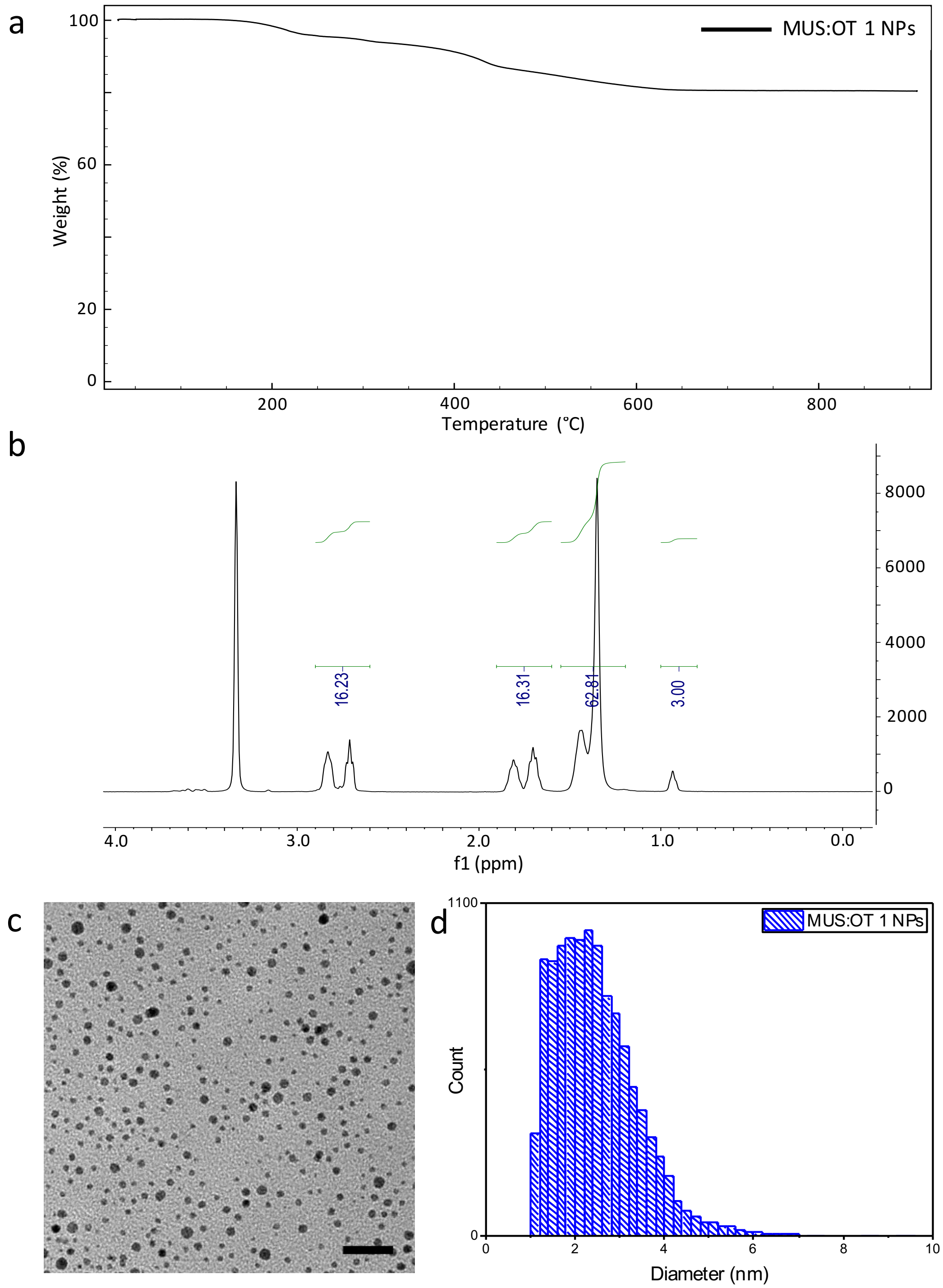

**Fig. S1**. Characterization of NPs (MUS:OT A as an example). a) Thermogravimetric analysis (TGA) plot of MUS:OT A NPs. The TGA curve is indicative of drying, desorption and thermal degradation of organic ligands. In the case of the MUS:OT NPs, OT desorbs between 176˚C and 233˚C, while MUS is decomposed at approximately 800˚C. The remaining weight at higher temperatures corresponds to the gold core of the NPs. The mass difference between the beginning and the end of this analysis allows for the ligand density estimation. b) 1H-NMR analysis of MUS:OT A etched with iodine solution in MeOD-d4 reveals the MUS:OT stoichiometric ratio, which in the case of MUS:OT A is 21 to 79, MUS to OT. c) Cryo-EM images of MUS:OT A. Scale bar is 20 nm. d) Particle size distribution of MUS:OT A obtained from the analysis of several cryo-EM images.

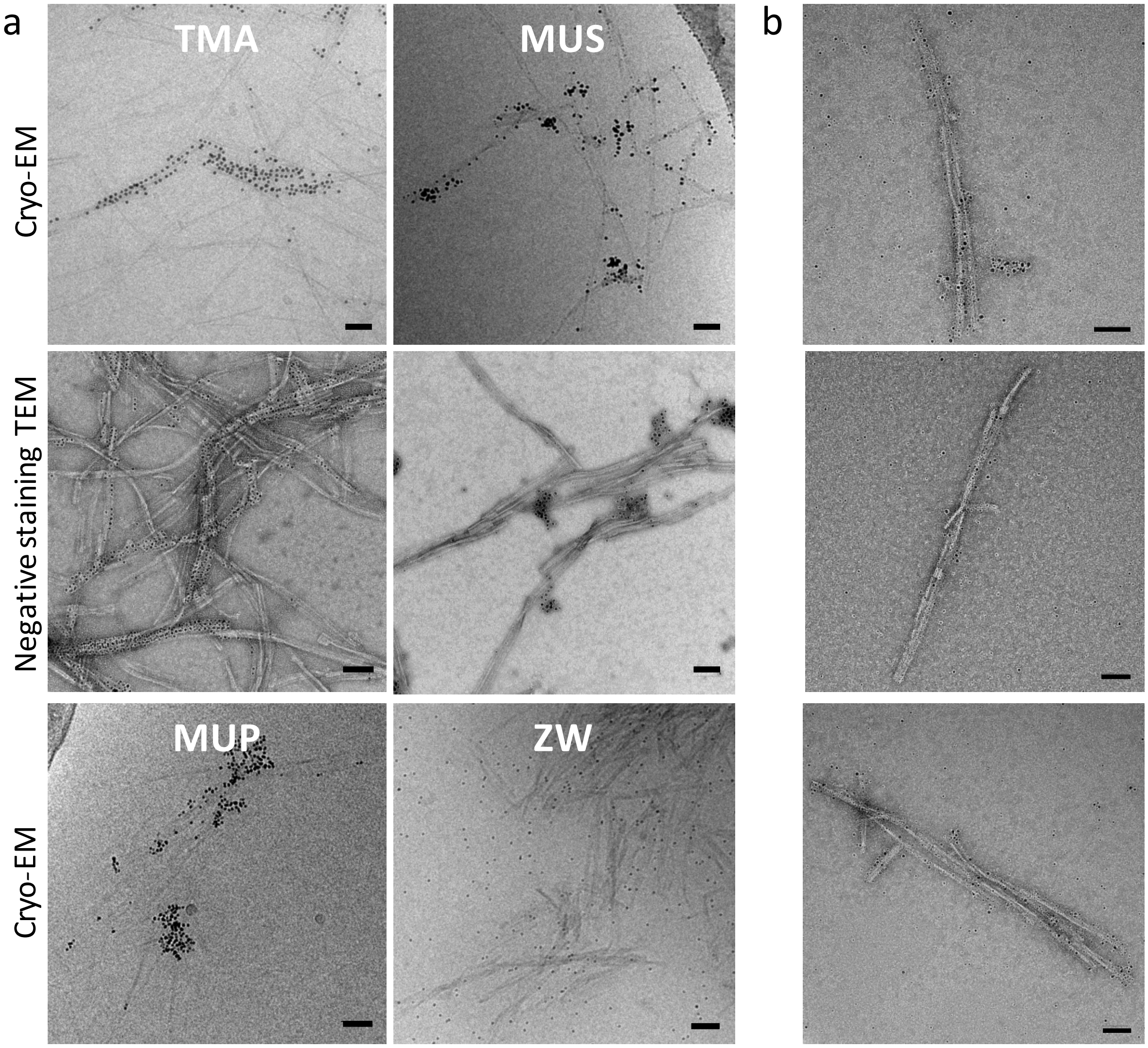

a
b
TMA
MUS
Cryo-EM
Negative staining TEM
MUP
ZW
Cryo-EM

**Fig. S2.** Decoration of $A\beta_{40}$ fibrils with various NPs. a) The upper panel shows cryo-EM images of $A\beta_{40}$ decorated with TMA and MUS NPs with the corresponding negative stain TEM images below. The bottom panel shows cryo-EM images of $A\beta_{40}$ fibrils incubated with MUP and ZW NPs. Cationic TMA NPs bind densely to the $A\beta_{40}$ fibrils and cause bundle formation. Cryo-EM microscopy suggests that these NPs decorated the fibrils in a cooperative manner, whereby some fibrils became densely decorated, while others remained bare. NPs protected only with negative MUP ligands have a propensity to aggregate in the solution rather than to decorate the fibrils, while NPs protected only with negative MUS randomly attached to the amyloids. Previous studies have demonstrated that the ZW NPs do not form protein coronas (13). Therefore, these NPs served as a control to assess the level of nonspecific interactions between the gold NPs and amyloids. We observe that the ZW NPs do not interact with the fibrils, indicating that the gold NPs do not interact nonspecifically with amyloid fibrils and that the effect observed in the case of both ratios of MUS:OT NPs is due to the adequate ligand coverage on the gold shell. b) Negative stain TEM images of $A\beta_{40}$ fibrils decorated with MUS:OT A NPs. All Scale bars are 50 nm.

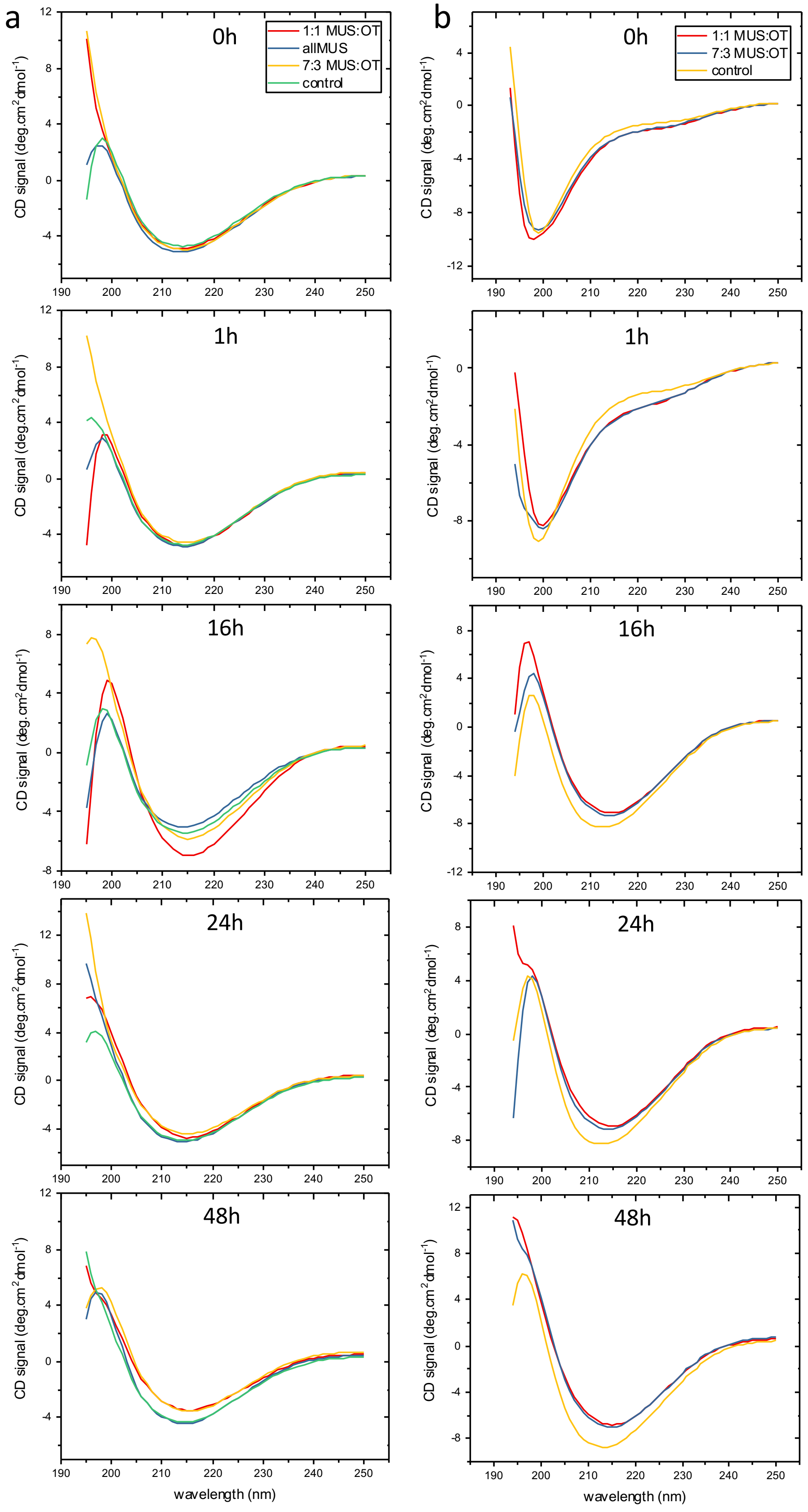
a
b
0h
1h
16h
24h
48h
1:1 MUS:OT
allMUS
7:3 MUS:OT
control
CD signal (deg.cm²dmol⁻¹)
wavelength (nm)

**Fig. S3**. CD spectra for A$\beta_{40}$ incubated with different NPs. a) CD spectra of mature A$\beta_{40}$ fibrils incubated with NPs. CD spectra were measured at different time points: 0 h, 1 h, 16 h, 24 h and 48 h. b) CD spectra of monomeric A$\beta_{40}$ incubated with NPs. CD spectra were measured at different time points: 0 h, 1 h, 16 h, 24 h and 48 h.

| Object [fibril] symmetry | | Fibril shape | Example of projection in cryo TEM | Referred in the paper as |
|---|---|---|---|---|
| Infinite | Twisted ribbon* | 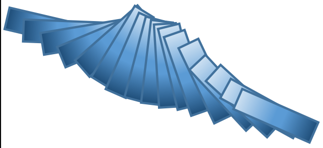 | 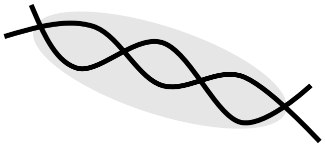 | Two fold |
| | Helical ribbon* | 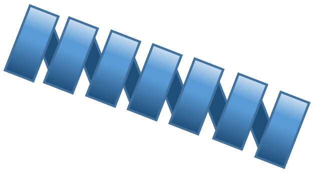 | 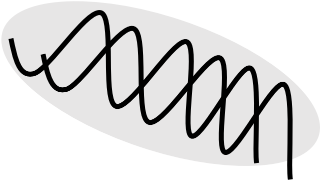 | Two fold |
| | Nanotube* | 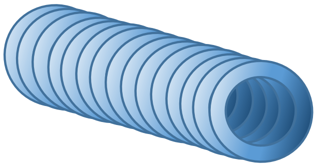 | 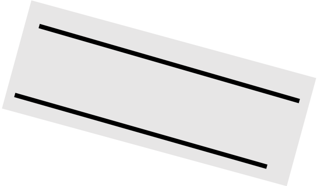 | Straight |
| Three fold | Three fold twisted fibril** | 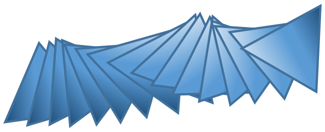 | 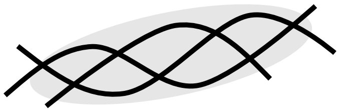 | Three fold |
| Two fold | Striated ribbon** | 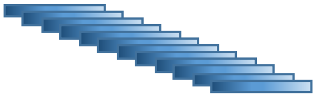 | 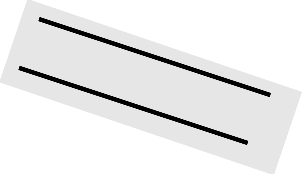 | Straight |

**Fig. S4**. Table showing possible amyloid shapes, symmetry and shape interpretations due to their 3D structure projected onto the 2D images. The structures noted with * were previously described by J. Adamcik and R. Mezzenga (14), and the structures marked with ** were described by Paravastu et al. (15).

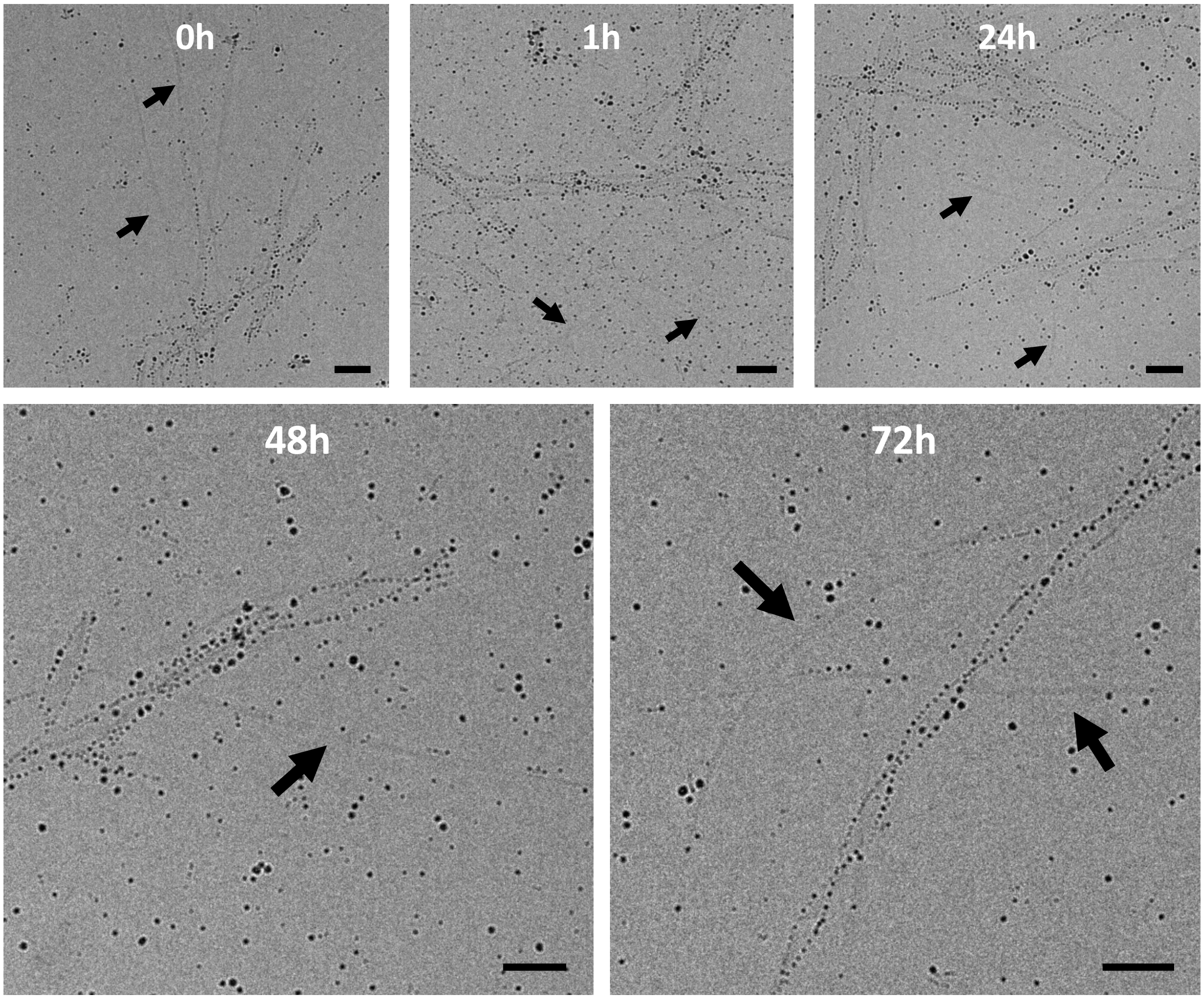


**Fig. S5.** R2 fibrils incubated with MUS:OT A NPs for various incubation times. The black arrows indicate bare structures still present in the solution after 3 days of continuous incubation. All scale bars are 50 nm.

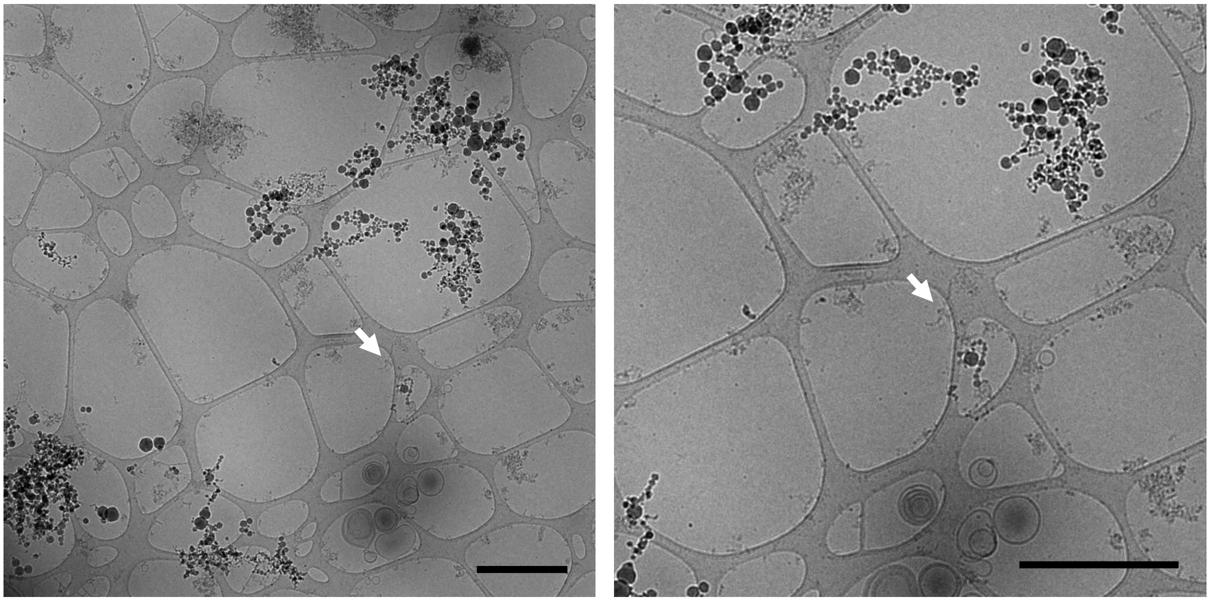

**Fig. S6.** Low-magnification images of the AD-derived PHF-enriched sample. The white arrow indicates PHFs. Scale bars are 1 µm.

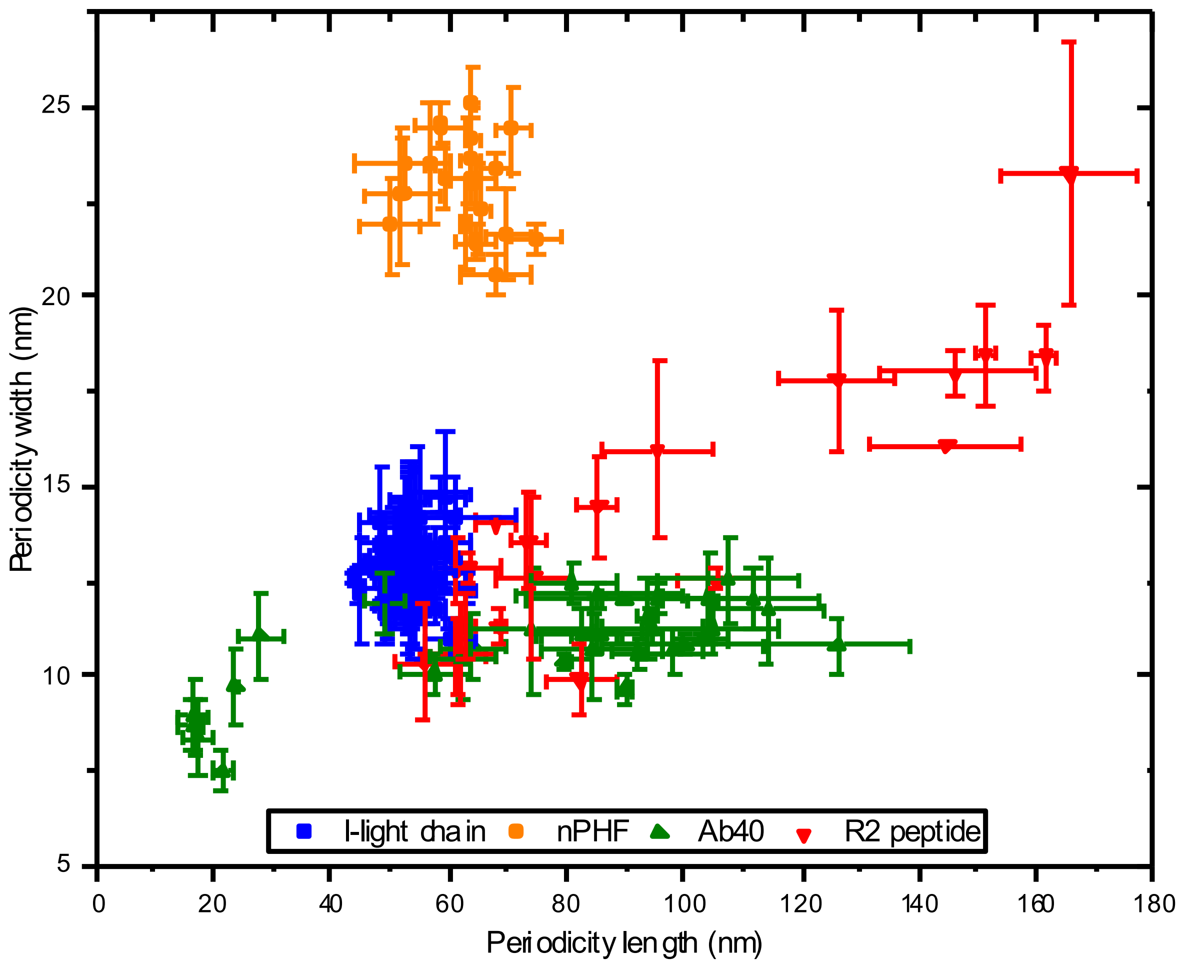


**Fig. S7.** Amyloid width plotted against crossover distance distribution with error bars. One dot represents the mean values obtained for the crossover distance and width of one fibril.

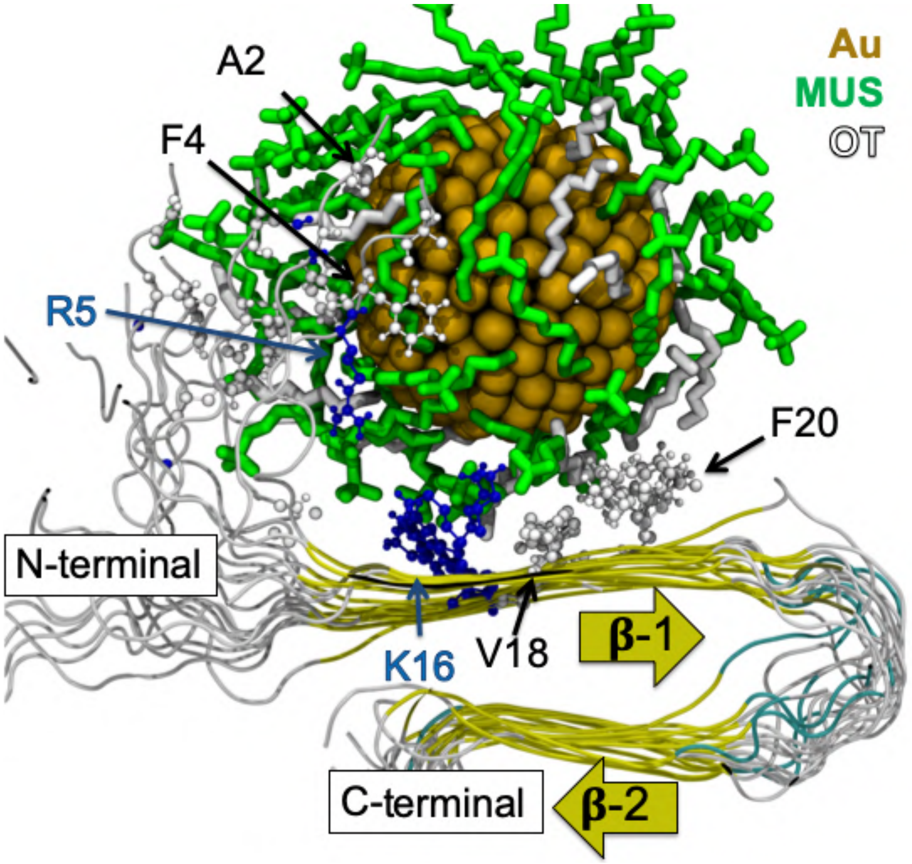


**Fig. S8.** Binding of the 7:3 NP to the A$\beta_{40}$ fibril. The protein is shown in cartoon representation and is colored and labeled accordingly to its secondary structure, while the amino acids that interact with the MUS and OT ligands are represented in ball-and-stick representation. Hydrophobic amino acids are shown in white, and positively charged amino acids are shown in blue.

### Supplementary Information – References


1. Uzun O, et al. (2008) Water-soluble amphiphilic gold nanoparticles with structured ligand shells. *Chem Commun* (2):196–198.

2. Zheng N, Fan J, Stucky GD (2006) One-step one-phase synthesis of monodisperse noble-metallic nanoparticles and their colloidal crystals. *J Am Chem Soc* 128(20):6550–6551.

3. Jan A, Hartley DM, Lashuel HA (2010) Preparation and characterization of toxic A aggregates for structural and functional studies in Alzheimer ' s disease research. *Nat Protoc* 5(6):1186–1209.

4. Fauvet B, et al. (2012) α-Synuclein in central nervous system and from erythrocytes, mammalian cells, and Escherichia coli exists predominantly as disordered monomer. *J Biol Chem* 287(19):15345–15364.

5. Harbison NW, Bhattacharya S, Eliezer D (2012) Assigning backbone NMR resonances for full length tau isoforms: Efficient compromise between manual assignments and reduced dimensionality. *PLoS One* 7(4):1–10.

6. Vieweg S, Ansaloni A, Wang Z-M, Warner JB, Lashuel HA (2016) An Intein-based Strategy for the Production of Tag-free Huntingtin Exon 1 Proteins enables New Insights into the PolyQ Dependence of Httex1 Aggregation and Fibril Formation. *J Biol Chem* 291(23):12074–12086.

7. Annamalai K, et al. (2016) Polymorphism of Amyloid Fibrils in Vivo. *Angew Chemie - Int Ed* 55(15):4822–4825.

8. Van Lehn RC, Alexander-Katz A (2013) Structure of mixed-monolayer-protected nanoparticles in aqueous salt solution from atomistic molecular dynamics simulations. *J Phys Chem C* 117(39):20104–20115.

9. Petkova AT, Yau W, Tycko R (2006) Experimental Constraints on Quaternary Structure in Alzheimer ' s -Amyloid. 498–512.

10. Buchete NV, Hummer G (2007) Structure and dynamics of parallel β-sheets, hydrophobic core, and loops in Alzheimer's Aβ fibrils. *Biophys J* 92(9):3032–3039.

11. Chalifour RJ, et al. (2003) Stereoselective interactions of peptide inhibitors with the β-amyloid peptide. *J Biol Chem* 278(37):34874–34881.

12. Tavanti F, Pedone A, Menziani MC (2018) Computational Insight into the Effect of Natural Compounds on the Destabilization of Preformed Amyloid-β(1–40) Fibrils. *Molecules* 23(6):1–15.

13. Moyano DF, et al. (2014) Fabrication of corona-free nanoparticles with tunable hydrophobicity. *ACS Nano* 8(7):6748–6755.

14. Adamcik J, Mezzenga R (2018) Amyloid Polymorphism in the Protein Folding and Aggregation Energy Landscape. *Angew Chemie - Int Ed* 57(28):8370–8382.

15. Paravastu AK, Leapman RD, Yau W, Tycko R (2008) Molecular structural basis for

polymorphism in Alzheimer ’ s β-amyloid fibrils. *Proc Natl Acad Sci U S A* 105(47):18349–18354.

**Author contributions**

P.J.S. and M.M. discovered the phenomenon, F.S., and H.A.L. designed the experiments and supervised the study, U.C. and P.J.S. designed the experiments and performed the research, N.Al B., S.V., A.Ch. and S.T. contributed to the preparation and characterization of the amyloid fibril preparations, M.M. contributed additional cryo-EM measurement, Z.P.G. contributed additional synthesis of nanoparticles and additional cryo-EM measurement, L.R. and M. F. contributed AL amyloid fibrils, F.T., M.C.M. and A. Al K. contributed molecular simulations. F.S., H.A.L., U.C., P.J.S., N.Al B., M.F. and L.R. contributed to the writing.

**Competing interests statement**

Authors declare no competing financial interests in association with this manuscript.
